# Proteogenomic Characterization of Primary Oral Squamous Cell Carcinomas Unveils the Extracellular Matrix Remodeling and Immunosuppressive Microenvironment Linked with Lymph Node Metastasis

**DOI:** 10.1101/2024.09.12.612653

**Authors:** Yu Liu, Zhenyu Yang, Jingya Jane Pu, Jie Zhong, Ui-Soon Khoo, Yu-Xiong Su, Gao Zhang

## Abstract

Oral squamous cell carcinoma (OSCC) is an increasingly prevalent malignancy worldwide. This study aims to understand molecular alterations associated with lymph node metastasis of OSCC in order to improve treatment strategies. We analyzed a cohort of 46 patients with primary OSCC, including 10 with lymph node metastasis and 36 without. Using a comprehensive multi-omics approach—encompassing genomic, transcriptomic, proteomic, epigenetic, single-cell, and spatial analyses—we integrated data to delineate the molecular landscape of OSCC in the context of lymph node metastasis. Our genomic analysis identified significant mutations in key genes within the MAPK, TGF-β, and WNT signaling pathways, which are essential for tumor development. The proteogenomic analysis highlighted pathways critical for lymph node dissemination and factors contributing to an immunosuppressive tumor microenvironment. Elevated levels of POSTN were found to reorganize the extracellular matrix (ECM), interact with TGF-β, disrupt cell cycle regulation, and suppress the immune response by reducing VCAM1 activity. Integrated analyses of single-cell and spatial transcriptome data revealed that cancer-associated fibroblasts (CAFs) secrete TGF-β1/2, promoting cancer cell metastasis through epithelial-mesenchymal transition (EMT). Our integrated multi-omics analysis provides a detailed understanding of molecular mechanisms driving lymph node metastasis of OSCC. These insights could lead to more precise diagnostics and targeted treatments.

## Introduction

Oral squamous cell carcinoma (OSCC) is the predominant oral malignancy, representing 80-90% of all oral cancers (1). The incidence of OSCC is rising, driven by factors such as smoking, alcohol consumption, betel quid chewing, human papillomavirus (HPV) infection, nutritional deficiencies, immune dysregulation, and genetic modifications (2, 3). These factors lead to genetic mutations, epigenetic changes, and an imbalanced microenvironment, contributing to the initiation and progression of OSCC (4). Clinically, OSCC often results in disfigurement and functional impairments, such as difficulties with swallowing, speech, and taste, significantly impacting patients’ quality of life (5). Moreover, OSCC is often diagnosed at advanced stages, posing challenges for treatment and management, with high recurrence rates and poor survival outcomes (6).

Despite advances in surgical and radiation therapies, the prognosis for OSCC patients remains poor due to the high incidence of lymph node metastasis (LNM) (7). Clinically, LNM is particularly challenging to detect in OSCC, as it may not be apparent in some cases and can only be detected through imaging or biopsy (6). Moreover, LNM drastically reduces the five-year survival rate from approximately 90% to about 40-50% and increases the likelihood of distant metastasis, particularly to the lungs, bones, and liver (8–10). Hence, identifying the molecular mechanisms and cellular transformations associated with LNM in OSCC is crucial for improving prognosis and reducing the risk of distant metastasis or tumor recurrence.

As a malignant epithelial tumor, OSCC presented highly metastatic potential due to the activation of the epithelial-mesenchymal transition (EMT) (11). EMT is a process during which cancer cells undergo phenotypic changes that include the loss of intercellular adhesion and apical-basal polarity (12). TGFβ is a key inducer of EMT, which leads to the disruption of epithelial-cell junctions, switches in cell elongation, and enhanced motility for directed migration and invasion through the extracellular matrix (ECM) (13). The alterations in the ECM can activate cell surface receptors (e.g., integrins) and initiate intracellular signaling cascades that promote EMT in cancer cells, ultimately contributing to cancer metastasis and invasion (14). Collectively, EMT, TGFβ pathway, and ECM remodeling are three essential factors collaborated for LNM in OSCC.

Numerous studies have explored biomarkers associated with OSCC progression through single-cell resolution, genomic profiling, and experimental analysis. Researchers have identified biomarkers such as *Cyclin L1*, *MMP10*, vimentin, *ROS 1*, *FGF8*, and *ZEB1*, and key pathways like NF-κB, ERK-STAT1/3, and EGFR (15–23). Additionally, pivotal cellular subtypes such as *CXCL8*-expressing cancer-associated fibroblasts, *LAIR2*-expressing Treg cells, and *SPP1+* macrophages have been implicated in OSCC metastasis (16, 24, 25). In addition, the dynamic immune microenvironment alterations may contribute to the development of LNM in OSCC, including an upregulation of PD-L1 expression on dendritic cells and an increase in the number of naive and quiescent CD4+ T cells, which have been linked to immune suppression (26, 27). Overall, these discoveries enhance our understanding of LNM in OSCC and provide a foundation for developing novel diagnostic and therapeutic strategies to improve patient outcomes.

However, several key scientific issues remain unresolved regarding LNM in OSCC progression. The influence of genomic mutations on pathways and biological processes facilitating cancer cell proliferation and metastasis is not fully understood. The transformation and aberrant crosstalk among the most critical cell types in the OSCC tumor microenvironment, including tumor cells, cancer-associated fibroblasts (CAFs), endothelial cells, and immune cells, have not been elucidated. Furthermore, the cooperation of genes and pathways with cellular communications in reconstructing the tumor microenvironment is unclear. Lastly, identifying key factors or cells that could be targeted for treating primary OSCC with LNM is essential. Therefore, a systematic exploration of the underlying mechanisms driving LNM in OSCC through comprehensive multi-omics research is urgent to develop more effective therapeutic strategies.

Currently, treatment for OSCC with LNM often involves a combination of surgery, radiation therapy, chemotherapy, and concurrent systemic therapy (targeted therapy and immunotherapy) (28). However, the optimal treatment approach is not always clear. It can depend on several factors, such as the number and location of metastatic lymph nodes, the extent of lymph node involvement, and the patient’s overall health. Furthermore, each of these therapeutic strategies exhibits restricted efficacy and outcomes due to the intratumoral and intercellular heterogeneity, constructing intricate cell interactions within the tumour microenvironment (TME) (29). Therefore, our study on the identification of key biomarkers associated with LNM in OSCC could potentially address some of these limitations and improve the treatment of this disease. By identifying patients who are at high risk of lymph node metastasis, our research could help clinicians make more informed treatment decisions and tailor treatment plans to individual patients. Additionally, by identifying new therapeutic targets and biomarkers, our study could lead to the development of more effective and targeted therapies for patients with LNM OSCC.

Next-generation sequencing (NGS) has significantly enhanced our understanding of the heterogeneity and evolution of different malignancies. Bulk tumor genomics, proteomics, and metabolomics offer paradigms for identifying disease-related biomarkers and potential molecular targets, advancing our comprehension of malignant transformation and therapeutic strategies (30–32). However, during tumor development and lymph node dissemination, OSCC exhibits aberrant cellular communications and tumor microenvironment remodeling, which traditional bulk tumor analyses cannot fully capture (33–35). Single- cell and spatial transcriptomics (ST) have emerged as powerful techniques for capturing transcriptomes with spatial resolution and monitoring target cell attributes during disease development stages (36, 37).

In this study, we assembled a cohort of primary OSCC with and without lymph node metastasis (pLN+ and pLN-) and performed multi-omics, single-cell, and spatial analyses to delineate the mechanisms associated with LNM in OSCC. Our comprehensive multi-omics analysis illuminates the genomic landscape of pLN+ and pLN- OSCC and characterizes the pLN+ OSCC as an “immune-suppressive ” subtype. Proteogenomics unveiled tumor environment during LNM dissemination may be fostered by elevated POSTN, potentially inducing ECM reorganization that interacts with TGF-β and disrupts cell cycle regulation to suppress the immune response. Moreover, single-cell and spatial transcriptome analysis indicated that the cytokines in the TGF-β pathway may be induced by CAFs, which in turn activate this pathway and facilitate CAF and cancer cell communication for promoting cancer cell proliferation and metastasis. These insights enhance our understanding of OSCC progression and highlight potential targets for novel therapeutic strategies.

## Results

### Patient Cohorts and Multi-Omics Data Analysis

Our goal was to investigate the proteogenomic landscape of lymph node (LN) metastasis in primary OSCC. Toward that goal, we retrospectively assembled a cohort of 50 patients who underwent surgical resection of primary tumors and lymph node dissections between July 2016 and December 2019 at Queen Mary Hospital (Hong Kong). After rigorous quality control, we eventually included 46 patients in this study **(Supplementary Fig. 1A)**. Their detailed demographics and clinicopathological features were presented in **Supplementary Table 1**. This cohort was divided into two groups, including 10 patients with LNM (pLN+) and 36 without (pLN-). Gender distribution was relatively balanced (45% males in pLN+ vs. 54% males in pLN-). The majority were aged between 50 and 69 years old (6 in pLN+ vs. 20 in pLN), non-smokers (9 in pLN+ vs. 26 in pLN), and non-drinkers (8 in pLN+ vs. 30 in pLN). Predominantly, tumors originated from the upper and lower gingiva (50% both in pLN+ and pLN-) **(Table 1)**. Statistical analyses revealed no significant differences in most clinicopathological features between these two groups (p>0.05), except for primary tumor subsites (p<0.05). All patients with LNM were diagnosed in advanced stages (stage III/IV) (AJCC 8th edition), confirming that there is an association between late- stage presentation and poor prognosis (38). Kaplan-Meier survival analysis highlighted significantly better outcomes for patients without LNM (pLN-), which was also corroborated by the analysis of an independent TCGA cohort of 92 patients with OSCC, including 41 pLN- and 51 pLN+ **(Supplementary Fig. 2B-C)** (39).

**Table 1.** Clinicopathological characteristics of recruited patients.

| Features | Primary OSCC<br>with lymph node<br>metastases<br>(pLN+) (n=10,<br>22%) | Primary OSCC<br>without lymph<br>node metastases<br>(pLN-) (n=36,<br>78%) | Overall<br>Cohort<br>(n=46) | P-Value |
| --- | --- | --- | --- | --- |
| **Sex** |  |  |  |  |
| Male, n (%) | 4 (40%) | 20 (56%) | 24 (52%) | 0.5714 |
| Female, n (%) | 6 (60%) | 16 (44%) | 22 (48%) |  |
| **Age (years)** |  |  |  |  |
| <50, n (%) | 1 (10%) | 9 (25%) | 10 (22%) | 0.4 |
| 50-69, n (%) | 6 (60%) | 20 (56%) | 26 (57%) |  |
| ≥70, n (%) | 3 (30%) | 7 (19%) | 10 (22%) |  |
| **Smoking** |  |  |  |  |
| Yes, n (%) | 1 (10%) | 10 (28%) | 11 (24%) | 1 |
| No, n (%) | 9 (90%) | 26 (72%) | 35 (76%) |  |
| **Alcohol Intake** |  |  |  |  |
| Yes, n (%) | 2 (20%) | 6 (17%) | 8 (17%) | 0.4667 |
| No, n (%) | 8 (80%) | 30 (83%) | 38 (83%) |  |
| **Subsites** |  |  |  |  |
| Buccal mucosa, n (%) | 3 (30%) | 4 (11%) | 7 (15%) | 0.009524 |
| Oral tongue, n (%) | 2 (20%) | 10 (28%) | 12 (26%) |  |
| Upper and lower gingiva, n (%) | 5 (50%) | 18 (50%) | 23 (50%) |  |
| Floor of mouth, n (%) | 0 (0%) | 2 (6%) | 2 (4%) |  |
| Hard palate, n (%) | 0 (0%) | 1 (3%) | 1 (2%) |  |
| Retromolar trigone, n (%) | 0 (0%) | 1 (3%) | 1 (2%) |  |
| **Pathology Staging (AJCC 8<sup>th</sup>)** |  |  |  |  |
| I, n (%) | 0 (0%) | 8 (22%) | 8 (17%) | 1 |
| II, n (%) | 0 (0%) | 10 (28%) | 10 (22%) |  |
| III, n (%) | 1 (10%) | 7 (19%) | 8 (17%) |  |
| IV, n (%) | 9 (90%) | 11 (31%) | 20 (44%) |  |

We then conducted comprehensive proteogenomic profiling, including whole-exome sequencing (WES) (n=41), RNA sequencing (RNAseq) (n=33), 4D-microDIA quantitative proteomics (n=24), DNA methylation arrays (n=25), single-nuclei RNA sequencing (snRNAseq) (n=4), and spatial transcriptomics (n=5) **(Fig. 1A, Supplementary Fig. 1A** and **Supplementary Table 2**). We also included two clinical cases as illustrative examples, including a patient, QM06, who is a 67- year-old male with non-metastatic OSCC at the mandible, and a patient, QM59, who is an 83-year-old female with metastatic OSCC, also at the mandible. Both CT and subsequent histopathological analysis confirmed the absence and presence of LN metastasis for QM06 and QM59, respectively **(Fig. 1B** and **1C)**.

**Figure 1.**
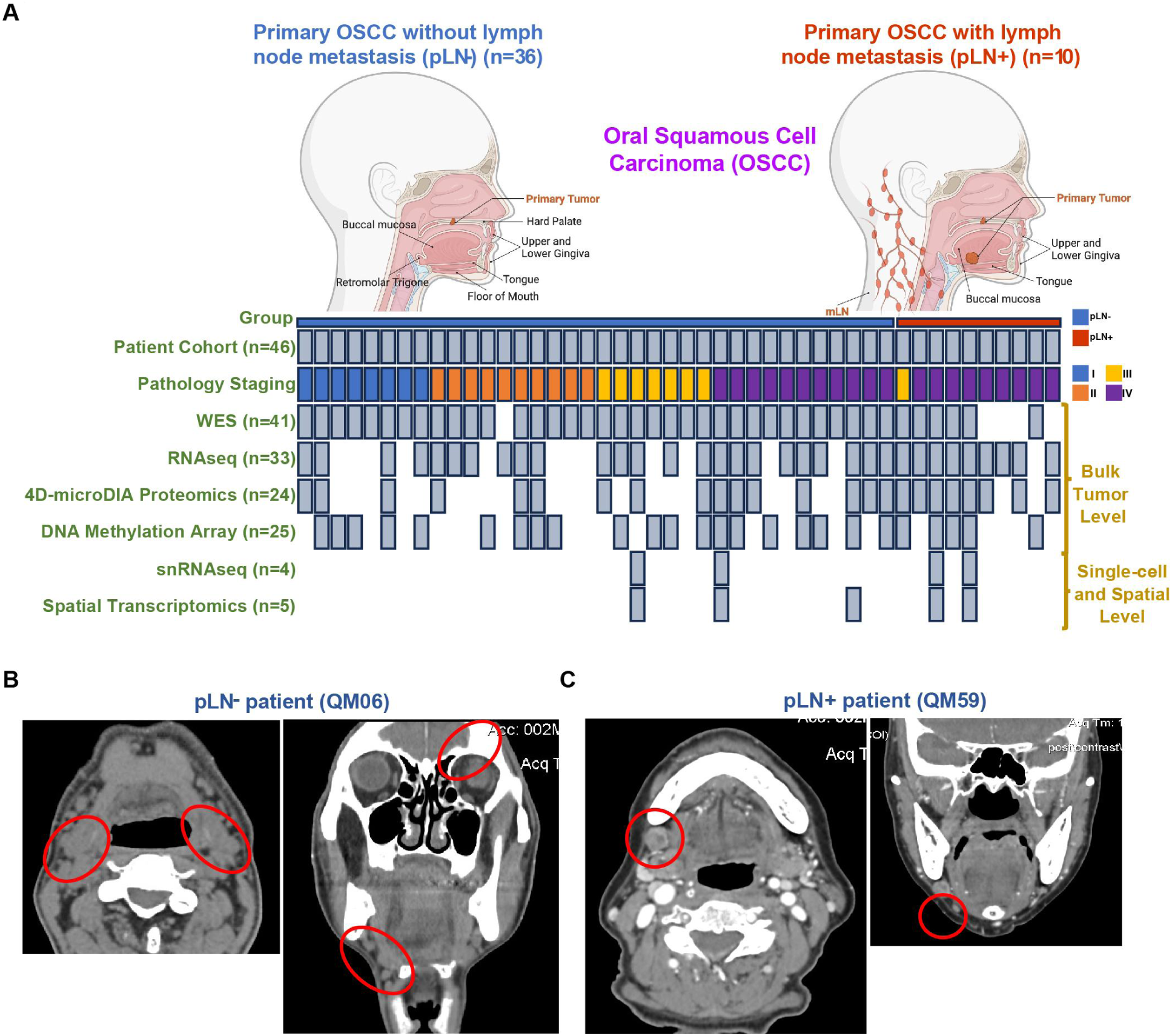
Patient cohort, overview of patients and samples, and experimental design. **A.** Schematic summary of two patient cohorts: pLN- (n=36, left) and pLN+ (n=10, right), along with the experimental platforms including WES, RNAseq, 4D-microDIA proteomics, DNA methylation array, snRNAseq, and spatial transcriptomics. Each column represents one patient’s tumor sample with profiling information annotated. **B.** Representative CT images from patients with squamous cell carcinoma at the mandible without lymph node metastasis in two views (left: axial, right: coronal). **C.** Representative CT images from patients with squamous cell carcinoma at the mandible with lymph node metastasis in two views (left: axial, right: coronal).

In summary, our study assembled a cohort of patients’ clinical samples that were suitable for a thorough molecular characterization of primary OSCC with and without LNM based on an integrative multi-omic approach.

### Genomic Insights into Primary OSCC with and without Lymph Node Metastasis

To establish the genomic landscape, we conducted whole-exome sequencing (WES) of primary OSCC samples without lymph node metastasis (pLN-) (n=35) and without lymph node metastasis (pLN+) (n=6). After processing WES data, we identified an average of 154 somatic mutations in pLN+ samples and 229 in pLN- samples by using Mutect2 (40). The tumor mutational burden (TMB) indicated a median of 2 mutations per megabase (Mb) for pLN+ and 3 for pLN-, respectively, encompassing single nucleotide polymorphisms (SNPs) and small insertions and deletions (Indels) **(Supplementary Fig. 2A)** (41). The variant allele frequency (VAF) analysis showed no significant intra-tumor heterogeneity (ITH) differences between the groups **(Supplementary Fig. 2B)** (42).

Our analysis highlighted alterations in *TP53, TTN, ANKRD36C, MUC5B, CDKN2A*, and *RETSAT* were shared by both groups, underscoring their roles in cell cycle regulation and tumorigenesis for primary OSCC in general (43–46). Unique to pLN- samples were *MUC16*, while *CPN1* and *KMT2A* were distinctive to pLN+ samples **(Fig. 2A-B)**. Subsequently, we illustrated the top 30 ranked mutations in both pLN- and pLN+ groups with regard to the mutational frequency of specific genes to further delineate the mutational profiles **(Fig. 2C-D)**. Specifically, we identified these mutated genes in the pLN- group were associated with multiple signaling pathways involved in tumorigenesis of HNSCC (*CSDM3, FAT1,* and *NOTCH1*), and cell cycle dysregulation (*CASP8, PRDM9,* and *SYNE1*) **(Fig. 2C)**. Likewise, the frequently mutated genes in the pLN+ group implicated the involvement of multiple signaling pathways which are related to tumorigenesis, including (1) WNT signaling pathway (*DCHS1*); (2) DNA repair signaling pathway (*FANCA*); (3) histone modifications signaling pathway (*KMT2A*); (4) TGF-β signaling pathway (*LTBP1*); and (5) MAPK signaling pathway (*MAP3K14, MAPK1,* and *MBP*) **(Fig. 2D)**.

**Figure 2.**
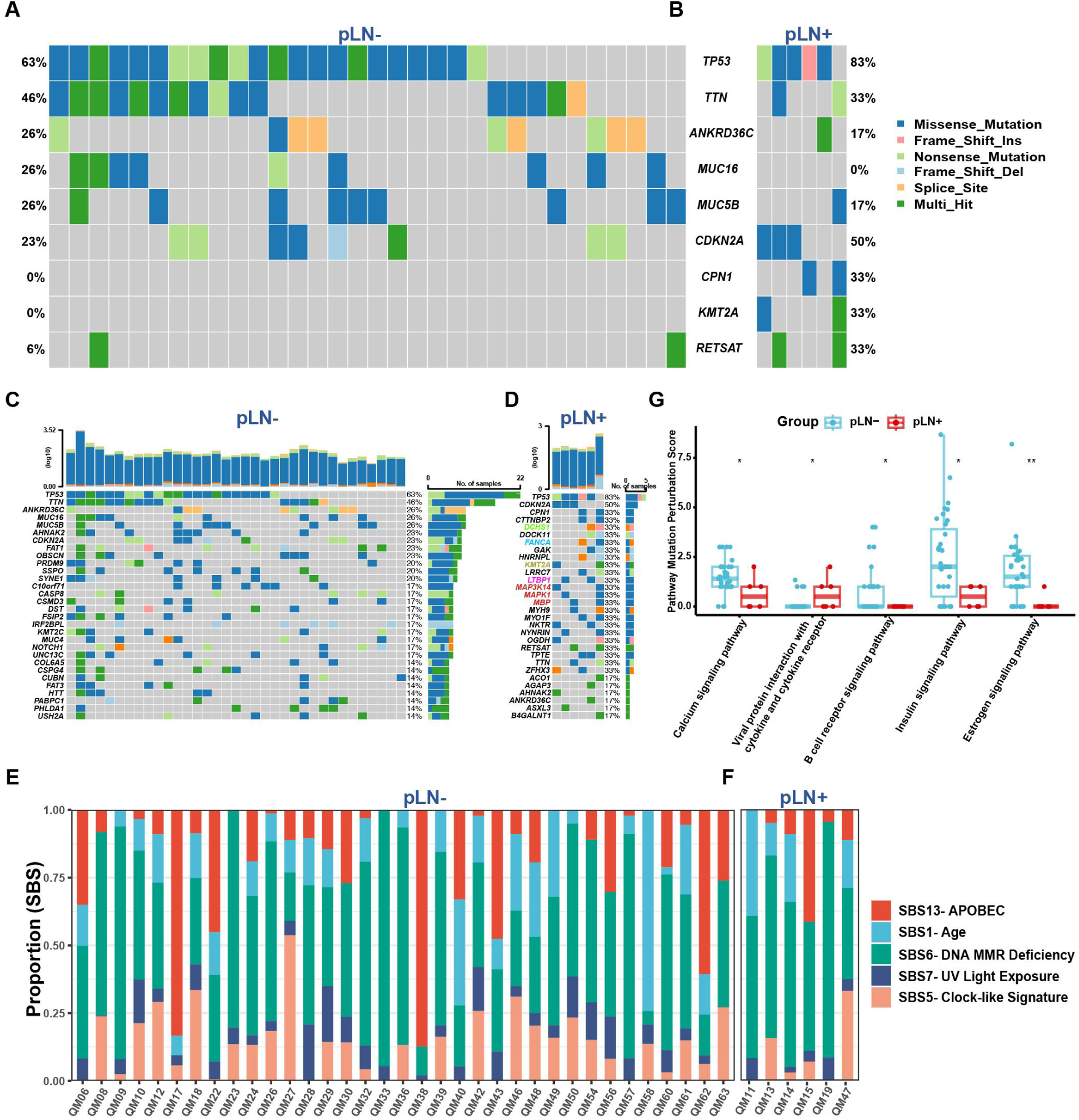
The genomic landscape of pLN- and pLN+ groups. **A and B.** Co-Oncoplots showing shared and unique mutated genes in pLN- **(A)** and pLN+ **(B)** tumor samples from the Queen Mary Hospital (QMH) cohort. **C and D.** Oncoplots showing individual mutated gene patterns in pLN- **(C)** and pLN+ **(D)** tumor samples. **E and F.** Percentage of mutational signature contribution for each tumor sample. **G.** Box plots showing pathway mutation perturbation (PMP) scores for each tumor sample.

We further extended our analysis to an independent cohort of 88 patients with OSCC (39 pLN- and 49 pLN+ OSCC) from TCGA to validate our previous findings, especially in *TP53, CDKN2A* and *TTN* **(Supplementary Fig. 2C-D)**. Not surprisingly, our analysis revealed a set of similar mutations of *CSMD3, FAT1, NOTCH1, CASP8, CDKN2A, SYNE1* and *MUC16* in the pLN- group (**Supplementary Fig. 2E**). Likewise, the analysis of the pLN+ group from the TCGA cohort unveiled two frequently mutated genes, *TP53* and *CDKN2A*, followed by *TTN*, *FAT1*, *NOTCH1*, *DNAH5* and *PCLO* **(Supplementary Fig. 2F)**.

The Catalogue of Somatic Mutations in Cancer (COSMIC) analysis revealed no significant differences between both groups with regard to key mutational signatures (47). Single base substitution (SBS) mutational signatures SBS 1 (age), SBS 5 (clock-like signature), SBS 6 (DNA MMR deficiency), SBS 7 (UV light exposure), and SBS 13 (APOBEC activity) were predominant signatures presented in most tumor specimens **(Fig. 2E-F and Supplementary Fig. 2G)**.

Next, we employed the Pathway Mutation Perturbation (PMP) score to assess the impact of mutations on specific pathways (48). Notably, the *‘Viral Protein Interaction with Cytokine and Cytokine Receptor’* pathway was significantly perturbed in the pLN+ group as compared to the pLN- group, suggesting activation or inhibition of cytokine signaling that possibly affects different aspects of immunity in cancer (49). Conversely, four pathways demonstrated significantly higher PMP scores in the pLN- group, including ‘*Calcium Signalling Pathway,’ ‘The B Cell Receptor Signalling Pathway,’ ‘Insulin Signalling* and *‘Estrogen Signalling Pathway’* **(Fig. 2G)**.

Taken together, our results delineated the genomic landscape and identified mutational profiles exhibited by primary OSCC with or without LNM, further offering insights into plausible molecular mechanisms of progression and metastasis of primary OSCC.

### Somatic Copy Number Alterations in Primary OSCC

To investigate somatic copy number alterations (SCNAs) and their implications for primary OSCC, we conducted the Genomic Identification of Significant Targets in Cancer (GISTIC) analysis (50). Specifically, the pLN+ group exhibited 17 arm-level losses, comprising 7 p-arms and 10 q-arms, and 18 peaks of deletion. In contrast, the pLN- group displayed 20 arm-level gains, including 9 p-arms and 11 q-arms, 23 peaks of amplification, and 18 peaks of deletion, indicating a more complex genomic alteration landscape **(Supplementary Fig. 3A-D)**.

Both groups shared all losses and deletions that we identified, with the most significant deletion peak at 18p11.32. Notable regions of alterations included amplifications at 3q29 (featuring genes *SDHA, RPL29, RPL35A*), 11q13.3 (*TIGAR*), and 11q22.2 (*MMP20*), and losses at 3p14.3 (*ADAMTS9*), 7q34 (*BRAF*), and 17q21.2 (*KRT13, CA9*) (51–61) **(Supplementary Fig. 3C-D)**.

The SCNAs exhibited by both primary OSCC groups elucidate a complex genomic landscape, highlighting the existence of specific genetic alterations that might contribute to the pathogenesis and metastatic potential of primary OSCC.

### The Analysis of Transcriptomic Data Reveals Key Insights into The Tumorigenesis of pLN+ OSCC

Next, we analyzed bulk RNA sequencing data derived from 33 tumor samples, including 9 pLN+ and 24 pLN- OSCC. The differentially expressed genes (DEGs) analysis identified that 7 genes were up- regulated and 173 were down-regulated in pLN+ tumors compared to pLN- (Log2fold Change > 1; adjusted Wald test p < 0.05) **(Fig. 3A and Supplementary Fig. 4A)**. Notably, *AGR2*, which significantly influences the EGFR signaling axis and tumor pathogenesis, exhibited the highest increase in the expression level (62, 63). Other up-regulated genes included *TGFβI*, known for its role in promoting metastasis through the TGF-β signaling axis and epithelial-mesenchymal transition (EMT), and *AURKA*, which is a key regulator of the cell cycle (64–66). Subsequently, we performed the Gene Set Enrichment Analysis (GSEA) by using an unbiased computational algorithm (Mann-Whitney-Wilcoxn Gene-Set Test, MWW-GST) based on ranked gene lists of DEGs from various gene set collections in the Molecular Signatures Database (MSigDB), including Gene Ontology Biological Process (GO-BP) (MSigDB C5), Kyoto Encyclopedia of Genes and Genomes (KEGG) (MSigDB C2), Hallmark (MSigDB H), and immunological signature gene sets (MSigDB C7) (67–69). Specifically, pLN+ OSCC showed enrichment of cell cycle pathways, particularly the G2M checkpoint and E2F targets, essential for cell proliferation and DNA replication (70, 71). In contrast, pLN- OSCC exhibited an enrichment of pathways related to lipid metabolism **(Fig. 3A)**. Therefore, to further validate these pathways enriched in the pLN+ OSCC, we interrogated another collection of gene sets, Reactome, and discovered 296 gene sets being significantly enriched (|normalized enrichment score| > 1 and adjusted p-value < 0.25) **(Supplementary Table. S3)**. This pathway analysis highlighted the involvement of cell cycle and DNA replication pathways in pLN+ OSCC, such as DNA strand elongation and telomere maintenance, alongside DNA repair pathways, including DNA unwinding and homologous recombination, that were critical for proliferation and metastasis of cancer cells **(Fig. 3B)**. Conversely, the signaling pathways enriched in the pLN- group, were predominantly associated with keratinization and muscle system processes. According to these findings, we carried out the functional analysis via aPEAR (Advanced Pathway Enrichment Analysis Representation) algorithm to identify the interconnected clusters based on the similarities between the pathway gene sets and construct a connective network (72). Notably, 6 clusters related to cell cycle and DNA repair were prominent in pLN+ OSCC, comprising (1) Gap filling DNA repair synthesis and ligation in GG-NER; (2) Homologous DNA pairing and strand exchange; (3) G0 and early G1; (4) Cyclin A/B1/B2 associated events during G2/M transition; (5) E2F-mediated regulation of DNA replication; and (6) Aberrant regulation of mitotic exit in cancer due to Rb1 defects. Furthermore, pathways involved in tumor invasiveness and metastasis were also connected and presented in pLN+ OSCC, including clusters named ‘MET promotes cell motility’ and ‘SMAD2/SMAD3:SMAD4 heterotrimer regulates transcription’. **(Fig. 3C)**. Taken together, our results underscore the enhanced proliferative and metastatic potential of pLN+ OSCC, thereby paving the avenue for further elucidating molecular mechanisms and cellular interactions exhibited by pLN+ and pLN- OSCC.

**Figure 3.**
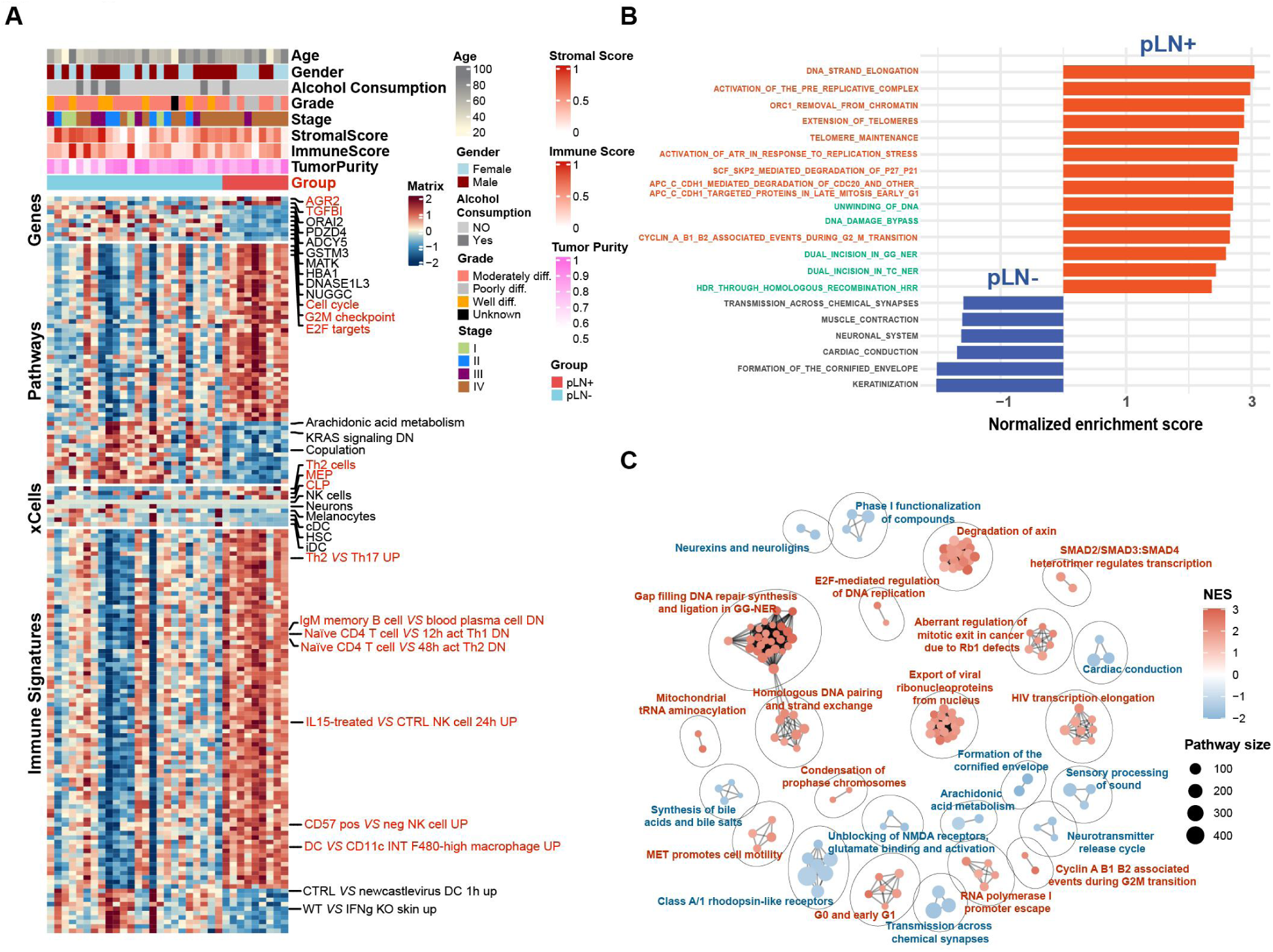
Transcriptomic characterization of pLN+ and pLN- groups. **A.** Heatmap illustrating two groups based on differentially expressed genes, enriched pathways, xCell- derived cells, and immune signatures. Upregulated genes, enriched pathways, abundant immune cells, and higher immune scores in pLN+ OSCC are highlighted in red. **B.** Gene Set Enrichment Analysis (GSEA) plots of the top 20 pathways (ranked by adj. P values) based on the Reactome collection from MSigDB. Red bars towards the right indicate enriched pathways in primary OSCC with lymph node metastasis; blue bars towards the left indicate enriched pathways in primary OSCC without lymph node metastasis. Pathways related to the cell cycle and DNA repair are highlighted in red and green, respectively. **C.** Network analysis presenting clusters across enriched pathways in the Reactome dataset based on similarity and connectivity. Clusters enriched in pLN+ OSCC are highlighted in red, and those enriched in pLN- OSCC are highlighted in blue.

Furthermore, to investigate the difference in immune infiltration between pLN+ and pLN- lesions, we conducted the CIBERSORTx (LM22) analysis of the bulk tumor RNAseq data to estimate the relative abundance of 22 immune cell populations in tumor samples from these two groups **(Supplementary Fig. 4B)** (73). The deconvolution analysis of immune cell populations demonstrated significant decreases in plasma cells within the pLN+ phenotype when comparing various cell subtypes. Therefore, to further evaluate the levels of immune cell infiltration, xCell and ESTIMATE immune scores were applied to assess the presence of different immune cells and to gauge immune scores in specific tumors (74, 75).

Our results demonstrated stronger signatures of T helper 2 cells (TH2), megakaryocyte–erythroid progenitor cells (MEP), and common lymphoid precursor (CLP) in pLN+ OSCC **(Fig. 3A)**. Notably, a prior study linked Th2 effector cells with unfavorable outcomes in OSCC due to the expression of *CCR8* (76). Correspondingly, via the enrichment analysis within immunological signature gene sets (MSigDB C7), the pLN+ phenotype showcased an upregulation of the Th2-involved pathway that is *Th2 VS Th 17 UP* **(Fig. 3A)**. Importantly, various studies have provided evidence supporting that activation of Th2 may contribute to the establishment of an immune-suppressive microenvironment, thereby facilitating tumor progression (77–80). In addition, other signatures related to immune suppression are also implied in the pLN+ OSCC, containing but not limited to *IgM memory B cell VS blood plasma cell DN*, *Naïve CD4 T cell VS 48h act Th2 DN*, *CD57 pos VS neg NK cell UP*. Thus, we speculated that pLN+ OSCC may adopt a phenotype of “immune-suppressive” compared with the pLN- OSCC.

To summarize, the analysis of transcriptomic data has revealed the presence of immune-suppressive signatures along with enrichment of the cell cycle and DNA repair pathway in the pLN+ phenotype, highlighting critical insights into the mechanisms underlying cancer cell dissemination.

### Characterization of The Proteome Network Identifies ECM Remodeling in pLN+ OSCC

We undertook the four-dimensional (4D) data-independent acquisition (DIA) quantitative proteomics approach to profile 24 fresh frozen tumor specimens, including 8 pLN+ and 16 pLN- (81). We subsequently identified 56 proteins that were significantly up-regulated in pLN+ OSCC, including TGFβI, which correlates with its mRNA elevation **(Supplementary Fig. 5A)**, RPS6KA4, PLAU and PTPN14, which were linked to key signaling pathways like MAPK, cell cycle and WNT(82–84).The protein enrichment analysis based on the Reactome collection of gene sets highlighted a significant enrichment of pathways related to extracellular matrix (ECM) remodeling in the pLN+ group, notably in “Extracellular Matrix Organization” and “Integrin Cell Surface Interactions” **(Supplementary Fig. 5B** and **Supplementary Table. S4)**. Our findings suggested that pLN+ OSCC exhibited an active ECM reorganization, which was conducive to tumor progression and metastasis (85).

Consistent with the aforementioned findings, we proposed to explore more biological characteristics from a proteome-wide perspective to investigate potential biomarkers of the dynamic alterations in pLN+ OSCC. To achieve this, we constructed a protein co-expression network based on protein expression profiling via weighted gene co-expression network analysis (WGCNA) (86). A scale-free network was constructed with scale-free R^2^ of 0.8 and soft-threshold power (β) of 12 as the soft-threshold values **(Supplementary Fig. 5C.D)**. Subsequently, by employing the “cutreeDynamic” function with minModuleSize = 20, we identified a total of 31 co-expression modules (ME) **(Supplementary Fig. 5E-F)**. Concurrently, we depicted the modules in a low-dimensional space and each was associated with unique biological processes and molecular pathways **(Fig. 4A** and **Supplementary Fig. 5F)**. These modules unveiled the essential processes during OSCC development and metastasis, encompassing ME1: Cytoplasmic translation; ME4: Cell substrate adhesion/Cell-matrix adhesion/; ME9: MYC targets v1; ME 10: G2M checkpoint; ME11: ECM receptor interaction/External encapsulating structure organization; ME20: Epithelial-mesenchymal transition; ME22:Immunoglobulin production; ME 26: Production of molecular mediator of immune response; and ME27: E2F target. Subsequently, by overlaying the abundance of differentially expressed proteins into the resulting WGCNA network (**Supplementary Fig. 5A**), we identified 6 modules (adj.P values < 0.05) of significant upregulation in pLN+ OSCC, including ME02, ME04 and ME11, whereas 3 modules exhibited significant elevation in pLN- OSCC, including ME07, ME15 and ME19 **(Fig. 4B)**. Specifically, all attributes featured in ME 11 were linked to ECM organization. Therefore, we designated ME11 as the ECM-related module **(Fig. 4A** and **Supplementary Fig. 5G)**. In addition, ME 07 was featured with dominant attributes associated with keratinization, which was collaborated with data derived from the transcriptomic analysis **(Fig. 4A** and **Supplementary Fig. 5H)**. Simultaneously, we calculated the degree centrality (DC) within each module and concentrated on the hub gene possessing top ranked DC scores. The 15 highest-ranked proteins in ME11 encompassed, but were not limited to, FN1, POSTN, LUM, COL6A2 and BGN, all of which were implied in ECM remodeling **(Fig. 4C)**. ME7 showcased the top 10 ranked hub proteins, among which CTNNB1 is a pivotal protein involved in the WNT and EGFR signaling pathways, which has been identified as a therapeutic target for tumorigenesis via a pan-cancer analysis **(Fig. 4D)** (87–89).

**Figure 4.**
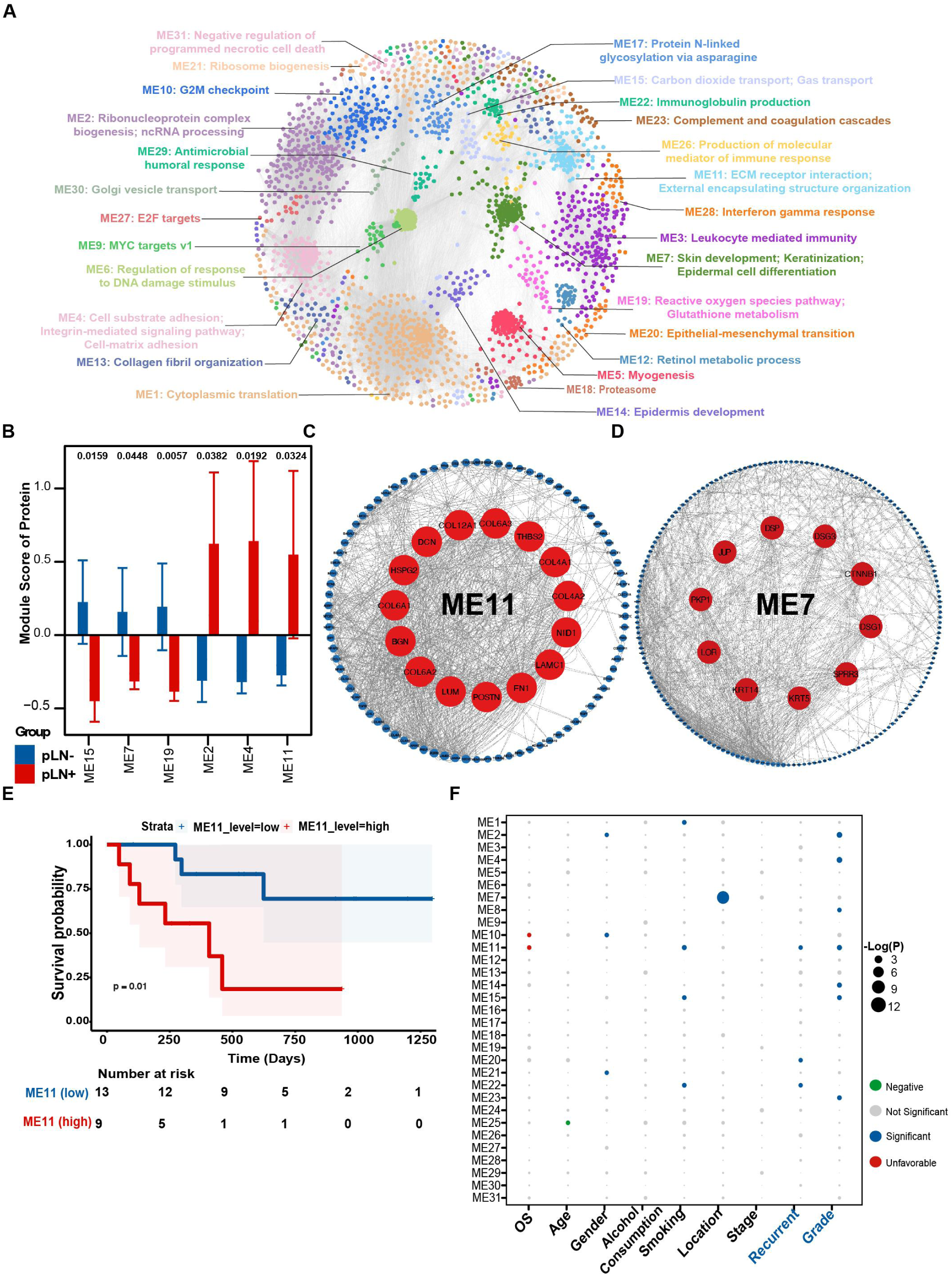
Protein co-expression network and clinical relevance of functional protein modules. **A.** WGCNA identified 30 functional protein modules (ME01–30) enriched in proteomic data for primary OSCC tumor samples. Each network node represents one protein, color-coded by different functional modules. **B.** Bar plot showing the normalized enrichment score with significant values of the 30 protein modules in pLN+ and pLN- OSCC. P values were calculated using the Mann–Whitney U-test and adjusted using the Benjamini–Hochberg method. Only modules with adjusted P < 0.05 are presented. Error bars represent means ± S.E. Two-sided P values were calculated. **C.** Sub-network of module 11. The top 15 ranked genes with the highest degree of centrality are highlighted in red and presented in the central circle. **D.** Sub-network of module 7. The top 10 ranked genes with the highest degree of centrality are highlighted in red and presented in the central circle. **E.** Kaplan–Meier survival curves comparing OS between patient subgroups stratified by high/low abundance (median cutoff) of ME11. P values were calculated using the log-rank test. **F.** Association of the enrichment score of the 30 modules with clinical data. Detailed information regarding the correlation analysis between module scores and clinical data is shown in Methods.

Subsequently, we conducted an association analysis to examine the clinical relevance of the identified modules. As expected, a high protein level of ME11 was found to be associated with an unfavorable prognosis of patients with LNM via the Kaplan-Meier analysis **(Fig. 4E)**. Furthermore, we calculated the enrichment score of each module with respect to its correlation with clinical data, such as overall survival (OS), age, gender, alcohol consumption, smoking, stage, recurrence, and grade. ME11 was found to be significantly enriched in characteristics that are essential for tumor recurrence and differentiated grade **(Fig. 4F)**. Additionally, ME11 was associated with poor OS, which is consistent with the results of the preceding Kaplan-Meier analysis.

Taken together, the analysis of 4D-DIA quantitative proteomics data not only strengthens the results of our transcriptomic data but also emphasizes ECM remodeling as a critical factor that underlies the progression and prognosis of pLN+ OSCC.

### Epigenetic Insights into Pathways Underlying Tumorigenesis of Primary OSCC

Epigenetic regulation is pivotal for the development of primary OSCC, influencing gene expression through mechanisms like DNA methylation and chromatin remodeling (90–92). To elucidate this, we performed a global DNA methylation analysis of 25 primary OSCC, including 20 pLN- and 5 pLN+ tumors by using the Infinium MethylationEPIC v2.0 BeadChip. This analysis targeted over 935,000 CpG sites, revealing 6,763 differentially methylated probes (DMPs) among all the samples. Subsequently, we interrogated the distribution of these DMPs within diverse functional genomic regions and CpG islands. Notably, changes were predominantly found within CpG islands, followed by N-shore, S-Shore, N-Shelf, and S-Shelf **(Supplementary Fig. 6A)**. Upon examing the sites of surrounding genes, we observed that differentially methylated sites were predominantly concentrated at the vicinity of gene bodies, followed by the intergenic region (IGR), transcription start sites (TSS1500, TSS200), 3’UTR, first exons, and 5’UTR **(Supplementary Fig. 6A)**. According to this, the CpG sites with differential methylation were subjected to consensus clustering analysis in order for us to visualize of the extent of DNA methylation between the two groups **(Supplementary Fig. 6B)**.

Among 117 differentially methylated regions (DMRs), a key finding was the demethylation of the *EGFR* gene promoter in pLN+ OSCC **(Supplementary Fig. 6C)**. Prior research has established that this modification likely facilitates an increase in *EGFR* transcription, strengthening the gene’s role in activating pathways crucial for the progression of primary OSCC (93).

Consequently, to elucidate the functional implications of these DNA methylation alterations, we performed the pathway analysis of genes with differentially methylated promoters or bodies. The analysis led to the identification of 64 Gene Ontology Biological Process (GOBP) terms, encompassing multicellular organismal process, multicellular organism development, and stem cell differentiation **(Supplementary Fig. 6D)**. Among these GOBP terms, notably, the top 5 ranked of them were (1) Multicellular organismal process; (2) Developmental process; (3) Anatomical structure development; (4) Multicellular organism development; (5) Cellular developmental process, which tend to be excessively activated during tumorigenesis of primary OSCC.

In summary, our epigenetic analysis not only identifies a significant demethylation of the *EGFR* promoter in pLN+ OSCC but also highlights the enrichment of pathways critical for tumor progression based on methylated regions. This analysis provides us with valuable insights into how primary OSCC were regulated epigenetically.

### Integrated Analyses of Multi-omics Data Highlights Key Biomarkers of pLN+ OSCC

When attempting the comprehensive data analysis, we conducted a correlative analysis of patient- matched mRNA and protein data. The results indicated that 81% out of 8,375 protein-coding genes exhibited a positive correlation, among which 26% of genes showed a statistical significance (mean Spearman coefficient of 0.23) **(Supplementary Fig. 7A)**.

We then chose the following criteria to select the potential prognostic proteins, including (1) a correlation coefficient greater than 0.7, (2) adjusted P value < 0.01, and (3) a hazard ratio (HR) greater than 2 for up-regulated proteins or HR less than 0.5 for down-regulated proteins **(Fig. 5A)**. Of the particular note, POSTN emerged as the protein with the highest positive correlation with prognosis, which was previously ranked as a top-ranked protein in the ME11 module **(Fig. 4C)**. POSTN (periostin), plays a pivotal role in tumor progression by interacting with ECM components and engaging WNT and NOTCH1 signaling pathways (94). Consequently, we further evaluated the survival data of the patients from our cohort based on protein and RNA expression levels of POSTN. Our analysis revealed that elevated expression of POSTN adversely correlates with survival for the patients with pLN+ OSCC in our cohort **(Fig. 5B-C)**.

**Figure 5.**
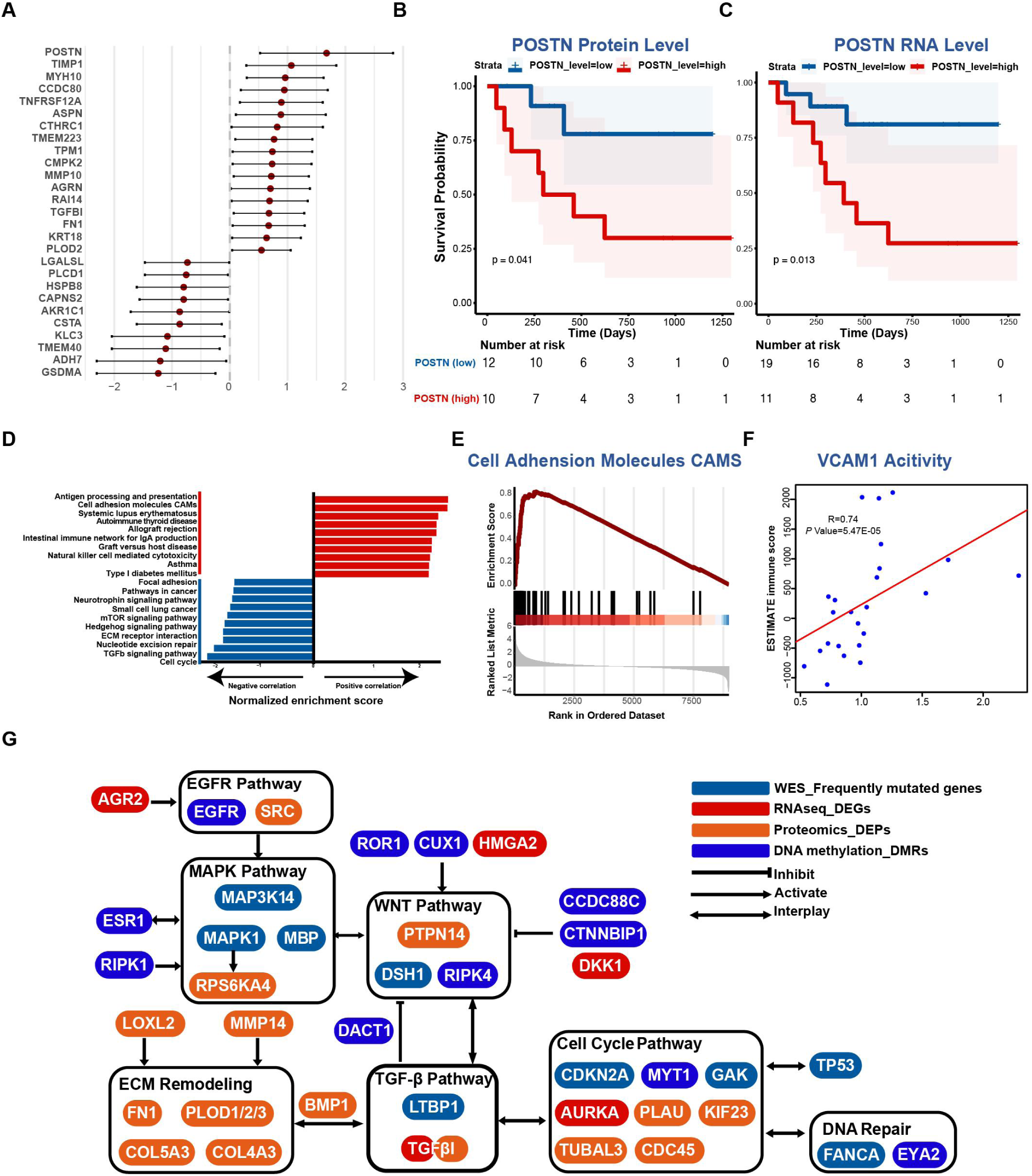
Integrative analysis of bulk tumor sequencing platforms. **A.** Forest plot showing the correlated effect of proteins screened from mRNA-protein correlation analysis. 27 proteins are presented with a correlation coefficient greater than 0.7, adjusted P value < 0.01, and hazard ratio (HR) greater than 2 or less than 0.5. **B and C.** Kaplan–Meier survival curves comparing OS between patient subgroups stratified by high/low expression (median cutoff) level of POSTN protein (B) and mRNA (C). P values were calculated using the log-rank test. **D.** Bar plot showing normalized enrichment scores for the top KEGG pathways correlated (red) or anti-correlated (blue) with immune scores. **E.** GSEA plot for cell adhesion molecules (CAMs) pathway correlated with immune scores. **F.** Scatterplot showing Spearman’s correlation of immune scores and VCAM1 activity. **G.** Molecular network showing alterations in pLN+ OSCC on genomic, transcriptomic, proteomic, and epigenetic levels. Genes identified in distinct platforms are highlighted in different colors.

Additionally, in order to delve deeper into immune features of pLN+ phenotype, we conducted the correlative analysis by incorporating ESTIMATE immune scores and mRNA-protein data **(Fig. 5D)** (74). As expected, immunological pathways demonstrated favorable correlations with these proteins, including antigen processing and presentation, autoimmune thyroid disease, intestinal immune network for IgA production, and natural killer cell-mediated cytotoxicity. However, signaling pathways like ECM receptor interaction, cell cycle, and TGF-β signaling showed negative correlations with immune scores, suggesting that their upregulation was linked with an immunosuppressive tumor microenvironment of pLN+ OSCC. Remarkably, the cell adhesion molecules (CAMs) pathway exhibited a pronounced positive correlation with immune scores, indicating the proteins in the CAMs family might be associated with immune infiltration **(Fig. 5D-E)**. Indeed, VCAM1 (Vascular Cell Adhesion Molecule 1, also referred to as CD106), which is an endothelial cell adhesion molecule, demonstrated a markedly positive correlation with immune infiltration scores **(Fig. 5F)**. Interestingly, the overexpression of TGF-β has been shown to suppress the expression of VCAM-1 in tumor endothelium, enabling tumor cells to evade immunosurveillance (95, 96). Collectively, our findings indicate that increases in cell cycle activity, TGF-β expression, and ECM remodeling may lead to the immune suppression by impeding the activation of VCAM1 expression.

Subsequently, we constructed a comprehensive molecular network that integrates genomic, transcriptomic, proteomic, and epigenetic data of pLN+ OSCC **(Fig. 5G)**. This integrated network revealed several pivotal signaling pathways pertinent to tumor development, encompassing EGFR, MAPK, WNT, cell cycle, DNA repair, TGF-β, and ECM-remodeling. Intriguingly, these pathways cooperate with each other to promote cancer cell invasion and metastasis, with several genes acting as the connections that work on two or more distinct pathways.

In conclusion, our integrated data analysis delineates a complex network of molecular interactions driving lymph node metastasis of pLN+ OSCC. Importantly, it highlights POSTN and VCAM1 as pivotal nodes influencing ECM remodeling and immune suppression, which interacts with cell cycle dysregulation and TGF-β expression to promot LN metastasis of primary OSCC.

### The Analysis of Single-cell RNA Sequencing and Spatial Transcriptome Data Reveals CAF- Secreted TGFβ1/2 Promotes Metastasis of pLN+ OSCC

Intra-tumor heterogeneity (ITH) and the tumor microenvironment (TME) are crucial in influencing the progression and metastasis of primary OSCC (97, 98). To explore this at both single cell and spatial levels, we leveraged advanced techniques of single-nuclei RNA sequencing (snRNAseq) and spatial transcriptomics. Specifically, four primary tumors were subject to snRNAseq, including 2 pLN+ and 2 pLN- OSCC (99, 100). After quality control, we identified 22,433 cells in total, which were further classified into 10 major clusters **(Supplementary Fig. 8A-B)**. By using markers of specific clusters, we identified five distinct cell types, including cancer-associated fibroblasts (CAFs) (*COL1A1*), endothelial cells (*PECAM1*), epithelial cells (*CDH1*), immune cells (*PTPRC*), and neurons (*RELN*) **(Fig. 6A-B)**. Among them, epithelial cells are commonly regarded as the origins of malignant cells that develop into the OSCC (33). In addition, since POSTN is an essential protein for activating the cell cycle/TGF-β/ECM-remoedling axis during LNM, we assessed the expression of POSTN and unveiled its higher expression in CAF than other cell types **(Supplementary Fig. 8C)**. Moreover, given that our integrated data analyses at the bulk tumor level revealed the up-regulation of the TGF-β signaling pathway, we focused on genes related to the TGF-β signaling when analyzing snRNAseq data. The analysis established that *TGF-β1* was highly expressed in CAFs and epithelial cells, whereas *TGF-β2* was mainly expressed in CAFs **(Fig. 6C-D)**. Collectively, these results not only provide additional evidence to support the involvement of POSTN protein and TGF-β signaling in the TME during LNM, but also pave the way for further investigations into the cellular interactions and diversity.

**Figure 6.**
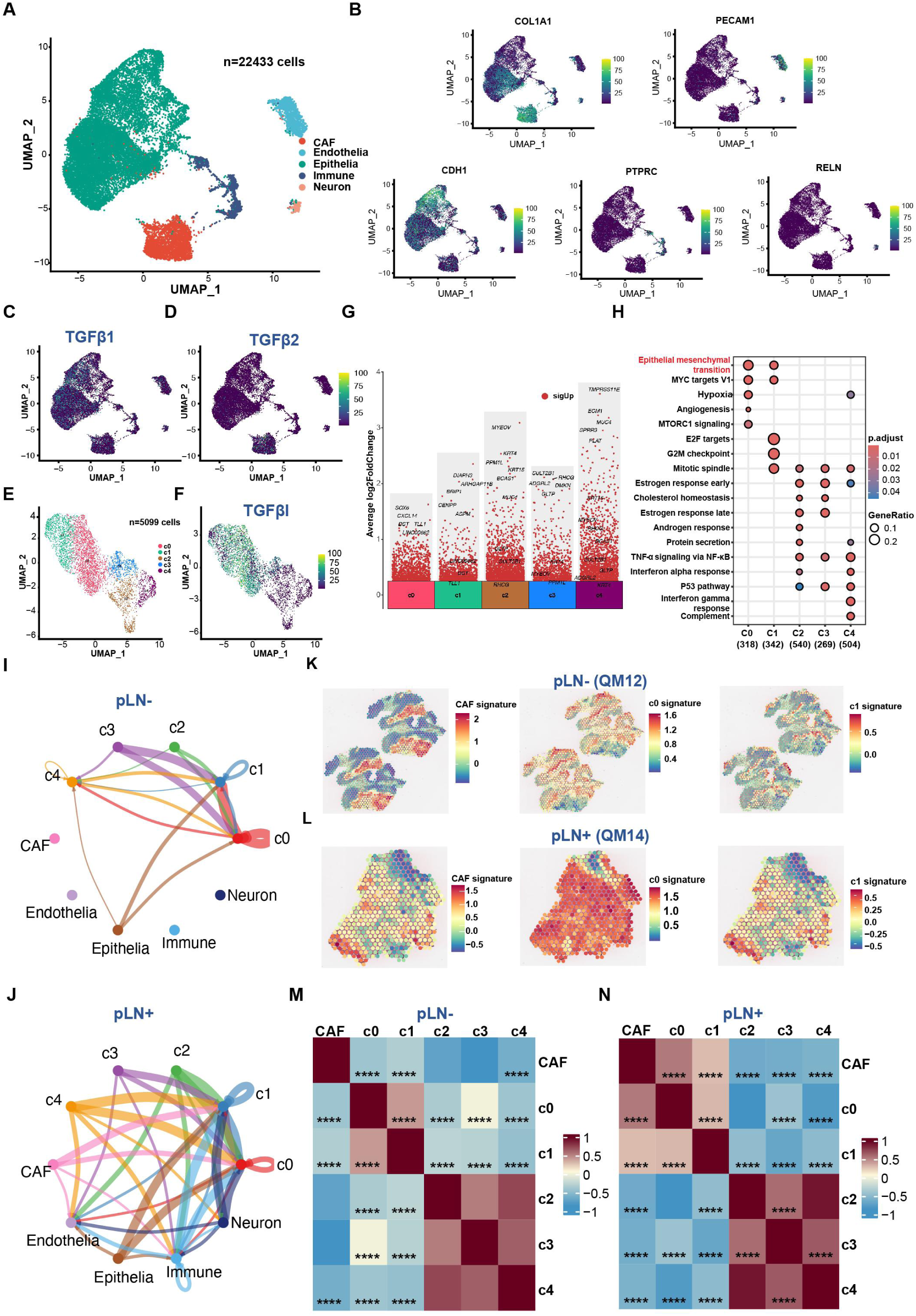
snRNAseq and spatial transcriptomic reveal crosstalk between CAF and cancer cells. **A.** UMAP plot of 22,433 cells derived from 4 OSCC patients showing 5 cell subclusters profiled by different colors. **B.** UMAP atlas showing the expression levels of 5 marker genes. **C and D.** UMAP atlas showing the expression levels of TGF-β related features, TGF-β1 **(C)**, TGF-β2 **(D)**. **E.** UMAP plot of 5,099 cancer cells scrutinized by CopyKat showing 5 cell subclusters profiled by different colors. **F.** UMAP atlas showing the expression levels of TGF-βI in cancer cells. **G.** Differential gene expression analysis showing up- and down-regulated genes across all 5 clusters. An adjusted p-value < 0.05 is indicated in red. **H.** Bubble plot showing enriched pathways across 5 clusters. The size of the dots represents the fraction of genes involved in the pathway, and the intensity of the color indicates the adjusted p-value. **I and J.** Circle plot showing intercellular TGF-β signaling interactions among different cell types and subclusters of cancer cells within pLN- **(I)** and pLN+ **(J)** OSCC. **K and L.** Spatial spots showing the expression level of CAF, C0, and C1 signatures in pLN- **(K)** and pLN+ OSCC **(L)**. **M and N.** Heatmap showing the correlation scores among different cell clusters in pLN- **(M)** and pLN+ OSCC **(N)**.

Subsequently, to further delve into tumor microenvironment presented in primary OSCC, we applied CopyKat (Copynumber Karyotyping of Tumors) to capture cancer cells (101). In total, we identified 5,099 cancer cells within the subset of epithelial cells, which were further clustered into five clusters (C0-C4) **(Supplementary Fig. 8D** and **Fig. 6E)**. pLN+ OSCC were primarily composed of C0 and C1 clusters, whereas pLN- OSCC were mainly consisted of C2, C3, and C4 clusters **(Supplementary Fig. 8E** and **Fig. 6E)**. Remarkably, the expression level of *TGFβI* was significantly higher in clusters C0 and C1 of pLN+ OSCC, designated as “TGFβI-positive” clusters **(Fig. 6F)**. Furthermore, to validate the cellular heterogeneity, we quantified cancer cell proportions in each cluster of pLN+ and pLN- OSCC. The data revealed that the “TGFβI-positive” cluster (C0 and C1) constituted over 80% of the pLN+ group, as compared to approximately 30% in the pLN- group. These results were consistent with our results based on the analysis of bulk tumor RNAseq data showing an increase in *TGFβI* expression in pLN+ OSCC **(Supplementary Fig. 8F)**. Subsequently, differential gene expression and gene set enrichment analyses showed genes related to cell-cell communication and ECM organization were over-expressed in the C0 cluster, such as *DST* and the enrichment of EMT pathways **(Fig. 6G-H)**. *DST*, *BRIP1,* and *CENPP* were up-regulated in the C1 cluster, leading to the enrichment of EMT and cell cycle-related pathways, including MYC targets, E2F targets, and G2M checkpoint **(Fig. 6G-H)**. Moreover, both clusters exhibited higher expression levels of TGF-β receptors *TGF-βR1* and *TGF-βR2*, indicating TGF-β signaling was activated in cancer cells of pLN+ OSCC **(Supplementary Table 5)**. Taken together, our results based on the activation of both upstream genes and downstream receptors may indicate the TGF-β signaling pathway was indeed activated in cancer cells within the pLN+ group.

Consequently, given that *TGF-β1* and *TGF-β2* genes are expressed in CAFs, we aimed to delineate cell-cell communication between CAFs and cancer cells since the cellular interaction might play a significant role in shaping the TME (102). Among these connections related to the TGFβ signaling pathway, we identified presence of a connection between the CAFs to C0 and C1 within the pLN+ OSCC, which is absent in the pLN- phenotype **(Fig. 6I-J)**. These results suggest that the activation of the TGF-β pathway in cancer cells may be induced by the cytokines (TGF-β1 and TGF-β2) that are secreted by CAFs.

To further complement these findings, we employed spatial transcriptomics (ST) of five fresh-frozen OSCC samples, including 2 pLN+ and 3 pLN- using the 10x Genomics Visium CytAssist platform. This approach preserved the 2D transcriptional information of cells, providing a spatial view of transcriptional heterogeneity within the TME (103, 104). Therefore, preceding to the snRNAseq findings, we subsequently picked up the top-50 ranked DEG in each subcluster of cancer cells, and evaluated the signature of CAF and each subcluster in both pLN+ and pLN- samples. Accordingly, the results illustrated the pLN+ OSCC exhibited elevated expression on the signature CAF, C0, and C1 clusters, compared to the pLN- OSCC **(Fig. 6K-L)**. Subsequently, in order to gain more information about the localization and interaction among these cells in the TME, we calculated the correlation analysis across these spatial spots. Not surprisingly, the pLN+ OSCC exhibited a significantly positive correlation among CAF, c0, and c1 clusters, whereas these correlations are significantly negative in pLN- OSCC **(Fig. 6M-N)**. These results suggest that CAF, C0, and C1 co-localize in the TME of pLN+ OSCC, which further validated the crosstalk between CAF and “TGFβI-positive” cancer cells. Furthermore, we performed Gene Set Variation Analysis (GSVA) to estimate the enrichment of gene sets in specific tumors, revealing that the EMT pathway was significantly enriched in the pLN+ sample in contrast to the pLN- sample **(Supplementary Fig. 8G-H)**. Collectively, these findings further validate that CAF could facilitate the invasiveness and metastasis of “TGFβI-positive” cancer cells via EMT during LNM in OSCC.

In summary, our single-cell and spatial transcriptome analyses reveal that CAF-secreted TGF-β1/2 plays a pivotal role in promoting metastasis in pLN+ OSCC by activating EMT pathways. These insights underscore the significance of TGF-β signaling in the TME and its potential to induce the lymph node dissemination in OSCC.

## Discussion

Oral squamous cell carcinoma (OSCC) remains a significant global health challenge, with over 370,000 new cases diagnosed annually (105). Early-stage OSCC is typically managed with surgery, while radiation therapy, either alone or combined with systemic therapy, is also commonly employed (106). However, for advanced stages involving metastasis, systemic therapies such as chemotherapy, targeted therapy, and immunotherapy become essential (107). Despite immunotherapy being a crucial component in the treatment of metastatic OSCC, its combination with curative radiotherapy or chemoradiotherapy has not consistently improved patient outcomes (108). Consequently, patients with lymph node metastases (LNM) face a significantly reduced prognosis and higher risks of distant metastases. Despite recent advances, the molecular mechanisms underlying early metastasis and the proteogenomic landscape of primary OSCC with LNM remain unclear, hindering the development of novel therapeutic targets. This study aimed to address this gap by conducting comprehensive multi-omics and spatial analyses of primary OSCC with or without LNM, to uncover the molecular underpinnings of cancer cell dissemination and inform treatment strategies.

Advancements in genetic sequencing technology have revolutionized precision oncology, enabling the identification of mutations and alterations in cancer genomes. However, sequencing data alone often fails to provide a complete understanding of the functional properties of genome-encoded proteins (109, 110). Quantitative proteomics has emerged as a promising approach to identifying disease biomarkers and molecular targets, enhancing our understanding of malignant transformation and therapeutic strategies (111–114). This study focused on integrated analyses of bulk tumor sequencing data, particularly from a proteomic perspective, an area previously underexplored in OSCC studies. Our proteomic layer-based multi-omics analysis of primary OSCC provided complementary insights beyond current genomic knowledge.

The extracellular matrix (ECM) plays a crucial role in maintaining tissue homeostasis, and its dysregulation is a hallmark of cancer progression (115). Our study revealed that ECM reorganization in LNM is driven by the activation of the transforming growth factor-β (TGF-β) signaling pathway and cell cycle dysregulation. This interaction contributes to an immunosuppressive tumor microenvironment, correlating with poor prognosis for LNM patients. Our findings are consistent with previous studies showing that TGF-β1-associated ECM genes recruit cancer-associated fibroblasts (CAFs), conferring immune evasion and resistance to cancer immunotherapies (116). Therefore, our study highlights the significance of ECM reorganization, TGF-β signaling activation, and cell cycle dysregulation in shaping the immunosuppressive microenvironment of LNM, presenting potential targets for novel therapeutic strategies.

*TGF-βI* (Transforming growth factor β induced) is a gene responsive to the TGF-β signaling pathway, induced by TGF-β1/2, and localized to the ECM (117). Our transcriptomic and proteomic analyses revealed its upregulation in pLN+ OSCC, with enrichment in the TGF-β and ECM remodeling pathways. Wang et al. found *TGF-βI* elevation in late-stage OSCC, indicating poor prognosis and potentially leading to an imbalanced immune response and OSCC progression (118). Additionally, *TGF-βI* has been shown to enhance the efficacy of chemotherapy drugs like paclitaxel, cisplatin, and gemcitabine in lung and ovarian cancers (119, 120). Thus, *TGF-βI* could serve as a novel biomarker for treating OSCC with LNM.

The immune system plays a critical role in defending against cancer, making immunotherapy a rapidly evolving field. However, lymph node metastasis impairs anti-tumor immunity, leading to immune suppression (121, 122). Single-cell studies in head and neck squamous cell carcinoma (HNSCC) patients have shown a decrease in stem-like CD8+ T cells and an increase in exhausted CD8+ T cells during LNM (27). Spatial transcriptomic analysis revealed that *CCXL12* expression leads to regulatory T-cell infiltration and increased TGF-β secretion, promoting a tumor immunosuppressive microenvironment (123). Our study builds on these findings, demonstrating that immunosuppressive signatures are exclusively present in pLN+ OSCC, providing further evidence that LNM induces an immunosuppressive microenvironment. This highlights the urgent need for a systematic exploration of this microenvironment and biomarkers to develop novel immunotherapies for metastatic disease.

Cell cycle dysregulation, a hallmark of cancer, leads to uncontrolled cell growth and tumor formation (124). Our transcriptomic analysis identified alterations in G2/M checkpoint genes in primary OSCC with LNM, which were negatively correlated with immune infiltration. This suggests that cell cycle dysregulation contributes to immune suppression during metastasis. Evidence from genomic and transcriptomic profiling indicates a link between immune evasion and cell cycle activity in cancer cells. Genetic amplification of *cyclin D1* (a CDK4/6 regulator) and *CDK4*, critical for cell cycle transitions, correlates with reduced efficacy of immune checkpoint blockade therapies (125). CDK4/6 inhibitors have shown promise in clinical trials, suggesting that targeting cell cycle proteins may effectively treat metastatic OSCC (126). The G2/M checkpoint has also emerged as a potential target, with inhibitors showing promise in treating glioblastoma (127). Thus, cell cycle inhibitors could represent novel therapeutic targets for metastatic OSCC.

POSTN (periostin), a matricellular protein involved in cell-matrix interactions, is often overexpressed in cancers, promoting tumor growth, invasion, and metastasis (128). POSTN activates PI3K/Akt and MAPK/ERK signaling pathways, regulates cell growth and metastasis, and shapes the tumor microenvironment (129, 130). It also modulates the immune response by stimulating immune cell recruitment and activation, facilitating tumorigenesis and invasion (131, 132). Our proteomic analysis identified POSTN as a pivotal protein in ECM remodeling, with elevated levels correlating with poor prognosis in LNM patients. Therefore, targeting POSTN could offer a promising therapeutic approach for OSCC patients.

Cancer-associated fibroblasts (CAFs) play various roles in OSCC, including promoting cancer cell proliferation, migration, invasion, and epithelial-mesenchymal transition (EMT) (133). Previous studies have shown that cancer cell-secreted molecules drive CAF activation in HNSCC, resulting in phenotypic changes and altered ECM production (134). Our single-cell and spatial analyses identified that CAFs secrete TGF-β1/2, enhancing TGF-βR1/2 expression in cancer cell subclusters in pLN+ OSCC. This suggests that CAFs activate the TGF-β pathway in cancer cells, facilitating communication and transformation between CAFs and cancer cells, which is exclusive to the pLN+ subtype. Therefore, targeting CAFs or their associated pathways could offer novel therapeutic interventions.

Given that cancer cell clusters in pLN+ OSCC are “*TGF-βI* positive” with enriched EMT functions, we speculate that TGF-βI is involved in upregulating EMT by inducing the marker genes in these subclusters, such as COL17A1, ITGA6, and CDH13. Future studies should examine the interplay between TGF-βI and these genes to better understand CAF and cancer cell crosstalk during OSCC metastasis. POSTN, identified in bulk proteogenomic analysis, may also play a critical role in these cell communications and microenvironment remodeling during cancer cell dissemination. Single-cell and spatial analyses have shown that *POSTN+*CAFs are enriched in advanced non-small cell lung cancer (NSCLC) tumors, associated with ECM remodeling, tumor invasion, and immune suppression (135). Therefore, we hypothesize that *POSTN+*CAFs are enriched in pLN+ OSCC and contribute to immune suppression, which warrants further exploration in future studies.

In summary, our integrated proteogenomic analysis has identified crucial pathways implicated in the tumorigenesis of primary OSCC with LNM and potential drivers of an “immune-suppressive” phenotype. This tumor microenvironment may be driven by augmented POSTN, ECM remodeling, TGF-β pathway activation, and cell cycle control disruption. In particular, the cytokines in the TGF-β pathway may be induced by CAFs, which in turn activate this pathway and facilitate CAF and cancer cell communication for promoting cancer cell proliferation and metastasis. These findings enhance our understanding of OSCC progression and highlight viable targets for developing novel therapeutic interventions. Further studies should explore the subtype of CAFs and identify key factors in the TGF-βI/EMT axis for OSCC progression and metastasis.

## Acknowledgments

The authors extend their gratitude to all collaborators for their insightful comments and discussions. This research was supported by the Oral Health Research and Innovation Fund (OHRIF) and the Faculty of Dentistry at the University of Hong Kong (G.Z.). Collaborative Research Fund (No. C7015-23GF), General Research Fund (No. 17117523), and Seed Fund for Collaborative Research (No. 2307102377), Hong Kong (Y.S.).

## Author contributions

G.Z. designed the study and provided financial and administrative support. Y.L. contributed to data interpretation and manuscript editing. G.Z. and Y.S. jointly supervised the study. J.P. provided study patients’ information and contributed to clinical data interpretation. Y.L. and Z.Y. contributed to data analysis and data interpretation. Y.L., J.Z., and U.K. participated in collecting clinical samples and data. All authors involved in writing the manuscript and final approval of the manuscript.

## Methods

### Biological Sample Collection and Processing

Tumor tissue and matched blood samples were collected from oral squamous cell carcinoma (OSCC) patients diagnosed at Oral and Maxillofacial Surgery Department, Queen Mary Hospital, Hong Kong, between Jul 2016 and Dec 2019. Patients consented to research use of their tissues, as approved by the Institutional Review Board of the University of Hong Kong/Hospital Authority Hong Kong West Cluster (IRB Reference Number: UW 15-239). Clinicopathological and follow-up data were retrospectively gathered from hospital records. Fresh tumor tissues were divided into two parts: one for formalin-fixed paraffin-embedded (FFPE) processing and the other quickly frozen for storage. Fresh frozen blood samples served as normal controls. Both fresh frozen tumor and blood samples were stored at −80°C until extraction, while FFPE samples were kept at room temperature for staining.

### DNA and RNA Extraction Germline DNA Extraction

Germline DNA was extracted from frozen blood samples using the TIANamp Genomic DNA kit (Catalog #: DP304) following the manufacturer’s instructions.

### Tumor DNA Extraction

Tumor DNA was extracted from fresh frozen tumor samples using Cetyltrimethylammonium Bromide (CTAB). The tissue block was ground with liquid nitrogen, and 50 mg was transferred to a 2.0 mL centrifuge tube containing 1 mL of CTAB lysis buffer. The mixture was incubated at 65°C with occasional mixing until fully lysed. The lysate was then centrifuged, and the supernatant was extracted with phenol (pH 8.0): chloroform: isoamyl alcohol (25:24:1), followed by chloroform: isoamyl alcohol (24:1). DNA was precipitated with isopropanol at -20°C, centrifuged, washed twice with 75% ethanol, and air-dried. The DNA was dissolved in ddH2O, incubated at 55-60°C if necessary, and treated with RNase A at 37°C for 15 minutes.

### Tumor RNA Extraction

Tumor RNA was extracted from fresh frozen tumor tissue using Trizol. The tissue block was ground with liquid nitrogen, and 50-100 mg was transferred to a 2.0 mL centrifuge tube containing 1 mL of Trizol. Chloroform (1/5 volume) was added, and the mixture was shaken vigorously, incubated at room temperature for 2-3 minutes, and centrifuged at 12,000 rpm at 4°C for 10-15 minutes. The upper aqueous phase was transferred to a new tube, mixed with an equal volume of isopropanol, and incubated at room temperature for 10 minutes. After centrifugation at 12,000 rpm at 4°C for 10 minutes, the supernatant was discarded, and the RNA pellet was washed with 75% ethanol prepared with DEPC water. The pellet was air-dried, dissolved in RNase-free water (DEPC water), and stored at -80°C.

### Whole-Exome Sequencing

Exome sequences were enriched from 0.4 μg of genomic DNA using the Agilent SureSelect Human All Exon V6 (Catalog #: 5190-8864) or Agilent SureSelectXT Mouse All Exon library (Catalog #: 5190-4643) according to the manufacturer’s protocol. Genomic DNA was fragmented to an average size of 180-280 bp using the Covaris S220. Overhangs were converted to blunt ends via exonuclease and polymerase activities. DNA fragments were end-repaired, phosphorylated, A-tailed, and ligated with paired-end adaptors at the 3’ ends. Adapter-ligated DNA fragments were enriched through PCR. Libraries were hybridized with biotin-labeled probes and captured using magnetic beads with streptomycin. Captured libraries were further enriched via PCR to add index tags for sequencing preparation. Libraries were purified using the AMPure XP system (Beverly, USA), analyzed for size distribution by the Agilent 5400 system, and quantified by QPCR (1.5 nM). Qualified libraries were pooled and sequenced on Illumina platforms using the PE150 strategy at CHI BIOTECH CO., LTD.

### RNA Sequencing

Total RNA was used to generate sequencing libraries using the NEB Next Ultra RNA Library Prep Kit for Illumina (Catalog #: E7530L). mRNA was purified from total RNA using poly-T oligo-attached magnetic beads. Fragmentation was performed with divalent cations under elevated temperatures. First-strand cDNA synthesis was conducted using random hexamer primers and M-MuLV Reverse Transcriptase (RNase H). Second-strand cDNA synthesis was performed using DNA Polymerase I and RNase H. Overhangs were converted to blunt ends, and NEB Next Adaptor with a hairpin loop structure was ligated to the adenylated 3’ ends of DNA fragments. Library fragments were purified to select 370-420 bp cDNA fragments using the AMPure XP system. USER Enzyme was used for size selection, followed by PCR with Phusion High-Fidelity DNA polymerase, Universal PCR primers, and Index (X) Primer. PCR products were purified, and library quality was assessed using the Agilent 5400 system and quantified by QPCR (1.5 nM). Qualified libraries were pooled and sequenced on Illumina platforms using the PE150 strategy at CHI BIOTECH CO., LTD.

### Computational Pipelines

All pipelines were developed according to National Cancer Institute sequencing guidelines. Tools from the GATK 4 suite were used for data processing.

### Alignment and Pre-processing

WES data pre-processing followed GATK Best Practices using GATK 4.0. Fastq files underwent quality control with ‘FastQC’. Sequencing adapters were removed with ‘Trimmomatic’. Fastq files were aligned to the hg38 genome using ‘BWA MEM’, and attributes were restored using ‘MergeBamAlignment’. PCR and optical duplicates were marked with ‘MarkDuplicates’. Base recalibration was performed using ‘BaseRecalibrator’ and ‘ApplyBQSR’. Coverage statistics were gathered using ‘CollectHsMetrics’. Alignment quality control was performed with ‘ValidateSamFile’ and inspected using ‘MultiQC’. ‘CrosscheckFingerprints’ confirmed readgroup integrity across samples, with mismatches excluded from further analysis.

RNA data pre-processing followed the mRNA analysis pipeline from the National Cancer Institute. Raw fastq files underwent quality control with ‘FastQC’. Bases failing quality control were trimmed using ‘Trimmomatic’. Fastq files were aligned to the hg38 genome using ‘STAR’ to generate BAM files. Post-alignment quality control was performed using ‘CollectRNASeqMetrics’. Missing values were imputed using the impute.knn package, with imputation on the genes present in at least 50% of samples.

### WES Variant Detection

Variant detection followed GATK Best Practices using GATK4. Germline variants were called from control samples using Mutect2 in artifact detection mode and pooled into a cohort-wide panel of normal samples. Somatic variants were called from tumor samples with matched normal controls using Mutect2, with parameters including the matched normal sample, reference fasta file, panel of normal, and gnomAD germline resources. Cross-sample contamination was evaluated using ‘GetPileupSummaries’ and ‘CalculateContamination’. Read orientation artifacts were assessed using ‘Collect-F1R2Counts’ and ‘LearnReadOrientationModel’. Additional filters were applied using ‘FilterMutectCalls’.

### WES Variant Post-Processing

BCFTools was used to normalize, sort, and index variants. A consensus VCF was generated, removing duplicate variants. The VCF file was annotated using GATK4.1 Funcotator (FUNCtional annOTATOR) with COSMIC, dbSNP, refGene data source.

### Mutational Burden

The mutational burden was calculated as the number of mutations per Mb sequenced. A minimum coverage threshold of 30x was required for each base. We regarded 38 Mb as the estimate of the exome size (136).

### Tumor Heterogeneity and MATH

Heterogeneity was inferred by clustering VAF in primary OSCC samples. The median absolute deviation (MAD) of mutant-allele fractions (MAF) was calculated for all tumors. MATH score, representing intratumor heterogeneity (ITH), was calculated as the percentage ratio of MAD to the median MAF value among the tumor’s mutated loci.

### Unique and Shared Mutations

Mutations in pLN+ samples were compared to matched pLN- samples. Shared mutations were defined as identical mutations at the same chromosomal positions leading to the same variants in the same genes. Unique mutations were those not shared between the groups.

### Mutational Signatures and Oncoplots

COSMIC mutational signatures were determined from mutations in primary OSCC samples. A mutation matrix was formed and decomposed into multiple signatures using NMF, compared to known COSMIC signatures based on cosine similarity. Differentially mutated genes were identified using a Fisher test and visualized with oncoplots.

### Copy Number Segmentation and Calling

Copy number identification was conducted via an open-source software called CNVkit with default settings, which is a tool kit to infer and visualize copy number from targeted DNA sequencing data (137). GISTIC2.0 identified SCNAs, with segmented copy numbers deconstructed using the ‘Ziggurat Deconstruction’ algorithm and significant SCNAs determined using the ‘Arbitrated Peel-off’ algorithm.

### Pathway Mutation Perturbation Level Measurement

Pathway activation was identified by mapping mutations to genes and applying criteria such as mutation frequency. Gene mutation perturbation scores (GMPscore) and pathway mutation perturbation scores (PMPscore) were calculated iteratively (48).

### Differential Gene Expression Analysis of RNAseq Data

Following alignment, BAM files were processed through the RNA Expression Workflow to determine RNA expression levels. Reads were mapped to each gene using Hisat2 (138). Transcripts were assembled using Stringtie, and the number of reads mapped to each gene was normalized using ‘DESeq2’, which employs a negative binomial distribution. DESeq2 provided base means across samples, log2 fold changes, standard errors, test statistics, p-values, and adjusted p-values. Significant genes were visualized using a ‘Volcano Plot’.

### Gene Set Enrichment Analysis

Pathway analysis was conducted using ‘fgsea’, a fast preranked GSEA. Ranked significant genes from ‘DESeq2’ and the Reactome pathway dataset (c2.cp.Reactome.v7.4) were used as inputs for ‘fgsea’, generating outputs including pathway names, enrichment scores, normalized enrichment scores, and p- values.

### Immune Cell Abundance Analysis

Relative immune cell fractions for downstream neoantigen analysis were determined using the ‘MCPcounter’ R package. ESTIMATE relative immune cell analysis was conducted using the ‘Estimate’ R package (74). Gene expression data was used in CIBERSORTx to estimate the abundances of member cell types in a mixed cell population.

### Mann-Whitney-Wilcoxon Gene-Set Test (MWW-GST)

To evaluate pathway activity, we calculated the normalized enrichment score of all pathways among four gene sets (GO-BP, KEGG, Hallmark, and immunologic signature gene set) using MWW-GST with the ranked list of DEGs. The MWW test statistic normalization provided the Normalized Enrichment Score (NES), an estimate of probability.

### Protein Extraction

Fresh-frozen tumor samples were ground with liquid nitrogen, and the powder was transferred to a 1.5 ml centrifuge tube. Samples were sonicated in a lysis buffer (8M urea with 1mM PMSF and 2mM EDTA), and debris was removed by centrifugation at 15,000g at 4°C for 10 min. Protein concentration was determined with a BCA kit. Equal amounts of proteins were digested with trypsin. The mix was reduced with 10 mM DTT for 45 minutes at 37°C and alkylated with 50 mM iodoacetamide for 15 minutes in the dark. Proteins were precipitated with chilled acetone at -20°C for 2 hours, air-dried, resuspended in 25 mM ammonium bicarbonate, and digested overnight at 37°C with trypsin. Peptides were desalted using a C18 Cartridge, dried, and redissolved in 0.1% formic acid.

### LC-MS/MS Detection

Liquid chromatography (LC) was performed on a nanoElute UHPLC. Approximately 200 ng peptides were separated over 60 minutes at 0.3 µL/min on a reverse-phase C18 column with an integrated CaptiveSpray Emitter. The temperature was maintained at 50°C. Mobile phases A and B were 0.1% formic acid in water and 0.1% formic acid in HPLC-grade acetonitrile (ACN), respectively. Mobile phase B was increased from 2% to 22% over 45 minutes, to 35% over the next 5 minutes, to 80% over the next 5 minutes, and held at 80% for 5 minutes. The LC was coupled online to a timsTOF Pro2 operated in Data- Dependent Parallel Accumulation-Serial Fragmentation (PASEF) mode with 10 PASEF MS/MS frames in one complete frame. The capillary voltage was set to 1400 V, and MS/MS spectra were acquired from 100 to 1700 m/z.

### MS Data Analysis

MS raw data were analyzed using DIA-NN (v1.8.1) with a library-free method. The Homo sapiens SwissProt database (20425 entries) was used to create a spectral library with neural network algorithms. MBR was employed to create and reanalyze a spectral library from DIA data. FDR was adjusted to < 1% at both protein and precursor ion levels. Identifications were used for further quantification analysis. Missing values were imputed using the impute.knn package, with imputation on the genes present in at least 50% of samples.

### Weighted Correlation Network Analysis (WGCNA)

Quantifiable proteins were analyzed with WGCNA to construct a protein co-expression network using the R package ‘WGCNA’ (86). Scale-free R^2^ = 0.8 was used for consistency with scale-free characteristics. The adjacency matrix was transformed into a topological overlap matrix (TOM) to reduce noise and spurious correlation. Network construction and module identification were based on TOM similarity. Parameters were set as follows: soft-threshold power (β) = 12, “cutreeDynamic” function, minModuleSize = 20. Functional annotation of each module was performed using the compareCluster subprogram of the R package ‘clusterProfiler’, DAVID, and STRING (139–141). Annotation gene sets for compareCluster analysis were downloaded from MSigDB (Hallmark gene sets, KEGG pathway database, and biological process from Gene Ontology) (139). The biological function of each module was summarized by the most significant enriched pathways (adjusted P < 0.05). Force-directed layout visualization of the 31 functional modules was created using the R package ‘igraph’. The protein co- expression network was visualized using Cytoscape v3.6.0 (142).

### Correlation Between Module Scores and Clinical Features

Statistical analysis of the correlation between module scores and clinical features included:

1. Prognosis Evaluation for OS Analysis: Patients were segregated into two groups based on the median module score. P values were calculated using the log-rank test. Modules were categorized as favorable (P < 0.05, HR < 1), unfavorable (P < 0.05, HR > 1), or not significant (P ≥ 0.05).
2. Continuous Factor Analysis: Spearman’s correlation explored relationships between module scores and continuous clinical features (e.g., age). Modules were classified as negative (P < 0.05, r < 0), positive (P < 0.05, r > 0), or not significant (P ≥ 0.05).
3. Categorical Factor Analysis: For binary factors (e.g., gender, alcohol consumption), P values were computed using the Mann–Whitney U-test. For factors with more than two categories (e.g., stage, grade), P values were derived using ANOVA. Significance was defined as P < 0.05.

### Survival Analysis

Kaplan-Meier analysis explored survival differences associated with lymph node metastasis group, ME expression level, and POSTN expression level. Statistical significance was assessed using Kaplan-Meier plots, log-rank tests, and Cox proportional hazards regression via the survminer (version 0.4.9) and survival (version 3.2-13) R packages. Patients who died for unrelated reasons were excluded.

### Illumina Infinium MethylationEPIC BeadChip and Data Processing

Genomic DNA concentration and integrity were assessed using a NanoDrop 2000 spectrophotometer and agarose gel electrophoresis. DNA was bisulfite-treated using the Zymo Research EZ DNA Methylation-Glod Kits. Bisulfite-converted DNA was analyzed on an Illumina Infinium MethylationEPIC v2.0 (935K) BeadChip and scanned using Illumina iSCAN. Idat files were preprocessed with the ChAMP (version 2.12.4) package in R and normalized using the BMIQ method (143). Statistical differences in continuous variables were compared by t-test.

### Differential Methylation Analysis

Differential methylation analysis was performed between 20 pLN+ and 5 pLN- tumor samples. Probes containing SNPs, probes in chromosome X, and probes with more than 10% missing values were excluded. The Wilcoxon rank-sum test determined differentially methylated CpGs (DMPs), with p-values adjusted by the FDR method. DMPs were reported if the mean methylation difference was >0.2 with an FDR of 5%.

### Differentially Methylated Region (DMR) Analysis

DMRs were identified separately for the two groups and combined using meta-analysis with the “dmrf” package in R and Rex (version 3.6.0) (144). Methylation levels of each CpG site were transformed using inverse normal transformation to ensure robustness against outliers and normal distribution assumptions. Regions with a maximum distance of 500 bp between consecutive features and at least two significant probes (p < 0.05) were identified. DMRs were evaluated using a Bonferroni adjusted significance level of 0.05. Annotation was performed using the ENSEMBL_MART_ENSEMBL BioMart database and the hsapiens_gene_ensembl database in the Ensembl genome browser (version: GRCh37).

### Gene Set Analysis (GSA)

Gene ontology (GO) terms were identified using significant CpGs and DMRs. Terms with at least five CpG sites were used to create a gene set, tested using a 0.05 FDR-adjusted significance level. GSA was performed using the “missMethyl” package in R (144, 145).

### mRNA-Protein Correlation Analysis

A total of 8,375 genes or proteins with less than 50% missing values were analyzed for gene-wise and sample-wise mRNA and protein correlations. Spearman’s correlation coefficient and corresponding P value were calculated for each mRNA-protein pair across pLN+ and pLN- tumors and individual samples using the cor.test function in R. Adjusted P values were calculated using the Benjamini-Hochberg (BH) procedure, with a cut-off of 0.01 for statistical significance.

### Prognostic Biomarker Analysis

To identify potential protein prognostic biomarkers, four criteria were used:

1. Proteins must be quantified in all samples.
2. The correlation coefficient between mRNA and protein expression should be > 0.7.
3. Candidate proteins should be differentially expressed between pLN+ and pLN- OSCC with adjusted P value < 0.01 (Wilcoxon signed-rank test, BH adjusted) and fold change > 1 at both mRNA and protein levels.
4. Kaplan-Meier curve with log-rank test visualized survival differences, and Cox proportional hazard model evaluated the hazard ratio (HR) for each protein. Candidate proteins should significantly correlate with overall survival (log-rank P value < 0.01, and HR (high/low) > 2 for upregulated or < 0.5 for downregulated proteins).

### Immune Scores Correlation Analysis

To identify potential drivers of immunosuppression, ESTIMATE immune scores were correlated with mRNA-protein data using Spearman’s correlation analysis. WebGestalt performed GSEA for KEGG pathways using signed -log10 P values (146). Gene set-based scores were the mean protein expression of all genes in that set.

### Single Nucleus Isolation and Sequencing

Nuclei were isolated using the Shbio Nuclei Isolation Kit (SHBIO, #52009-10, China) and counted with a cell counter (Thermo Fisher). Using a Chromium Single Cell 3′ Library and Gel Bead Kit v3 (10X Genomics), nuclei were loaded onto a Chromium Single Cell Processor (10X Genomics) to barcode RNA from single nuclei. Sequencing libraries were constructed according to the manufacturer’s instructions (10× Genomics) and sequenced on a NovaSeq 6000 system (Illumina, 20012866) (147).

### snRNA-seq Data Processing

Reads were processed using the Cell Ranger 3.0.1 pipeline with default and recommended parameters. FASTQs from Illumina sequencing output were aligned to the human genome, version GRCm38, using the STAR algorithm (148). Gene-Barcode matrices were generated by counting UMIs and filtering non- cell associated barcodes. The gene-barcode matrix containing barcoded cells and gene expression counts was imported into Seurat_4.1.3 R toolkit for quality control and downstream analysis of single-cell RNAseq data (149).

### snRNAseq Clustering Analysis

Cluster analysis of single-cell count matrices was performed using the R package ‘Seurat’ (v4.1.3) (150). Normalization and scaling were performed after filtering using the ‘NormalizationData’ and ‘ScaleData’ functions with default parameters. Principal components for highly variable genes were calculated using ‘RunPCA’. After quality control, removal of batch effects, and data integration from 4 samples (2 pLN+ and 2 pLN-), 22,433 cells were used in downstream analysis. Clusters were identified using ‘FindClusters’ with a 0.5 resolution. Uniform manifold approximation and projection (UMAP) visualized clusters in a reduced 2D space. Cluster markers were identified using ‘FindAllMarkers’, and cell types were assigned using cluster markers and CellMarker 2.0 (151).

### Cancer Cell Prediction

The CopyKAT package predicted cancer cells from epithelial cells (101). Cells from non-epithelia war regarded as reference, while cells with considerable aneuploidy mutations were considered cancer cells. Each sample was calculated individually and all cells with aneuploidy mutations were annotated for downstream analysis as a new cell type.

### Cell-Cell Communication Analysis

The R package Cellchat (version 2.1.2) inferred interaction mechanisms among tumor microenvironment (TME) components across pLN+ and pLN- tumor tissues (152). The ‘anadata’ dataset was transformed to generate a new Cellchat object. Functions such as ‘compareInteractions’, ‘netVisual_heatmap’, ‘netAnalysis_signalingRole_scatter’, and ‘rankNet’ analyzed and compared interaction numbers, strengths, and information flow of signaling pathways or ligand-receptor pairs among tissues. Interaction numbers and strengths among different cellular components within tissues were also investigated.

### Spatial Sequencing Library Preparation

Samples with RNA quality control (RIN > 4) were used for spatial transcriptomic construction and sequencing. 10-micron thick sections were mounted onto glass slides, fixed in ice-cold methanol, stained with hematoxylin and eosin, and scanned under a microscope (Keyence, Itasca, IL, USA). The stained slide was incubated with a Human whole transcriptome probe panel and transferred to Cytassist (10X Genomics). A human whole transcriptome probe panel (10x) consisting of three pairs of specific probes for most genes was hybridized to RNA. Probe pairs were ligated, forming single-stranded ligation products. Samples were treated with RNase and permeabilized to release ligation products. Poly-A portions of products were captured by poly(dT) regions of capture probes on the Visium slide, including an Illumina Read 1, spatial barcode, and unique molecular identifier (UMI). Probes were elongated to generate spatially barcoded ligated probe products, detached from the slide, indexed through Sample Index PCR, and sequenced. Visium Spatial Gene Expression libraries consisted of Illumina paired-end sequences flanked with P5/P7. The 16-bp Spatial Barcode and 12-bp UMI were encoded in Read 1, while Read 2S sequenced the ligated probe insert.

### Signature Expression Analysis of Spatial Spots

We selected the top 50 ranked DEGs in each cluster as the signatures, then calculated the cluster scores for feature expression programs in spatially single cell level via addmodulescore function in ‘Seurat’ R package (version 4.4.0), with the default settings.

### Gene Set Variation Analysis (GSVA)

Pathway activities of tumor cluster spots were quantified using GSVA, implemented in the GSVA package (153). The log-transformed normalization expression matrix of tumor spots was input into the “gsva” function with default parameters. A set of 50 cancer hallmark signatures was used for analysis.

## Figure and Table Legends

**Supplementary Figure 1.**
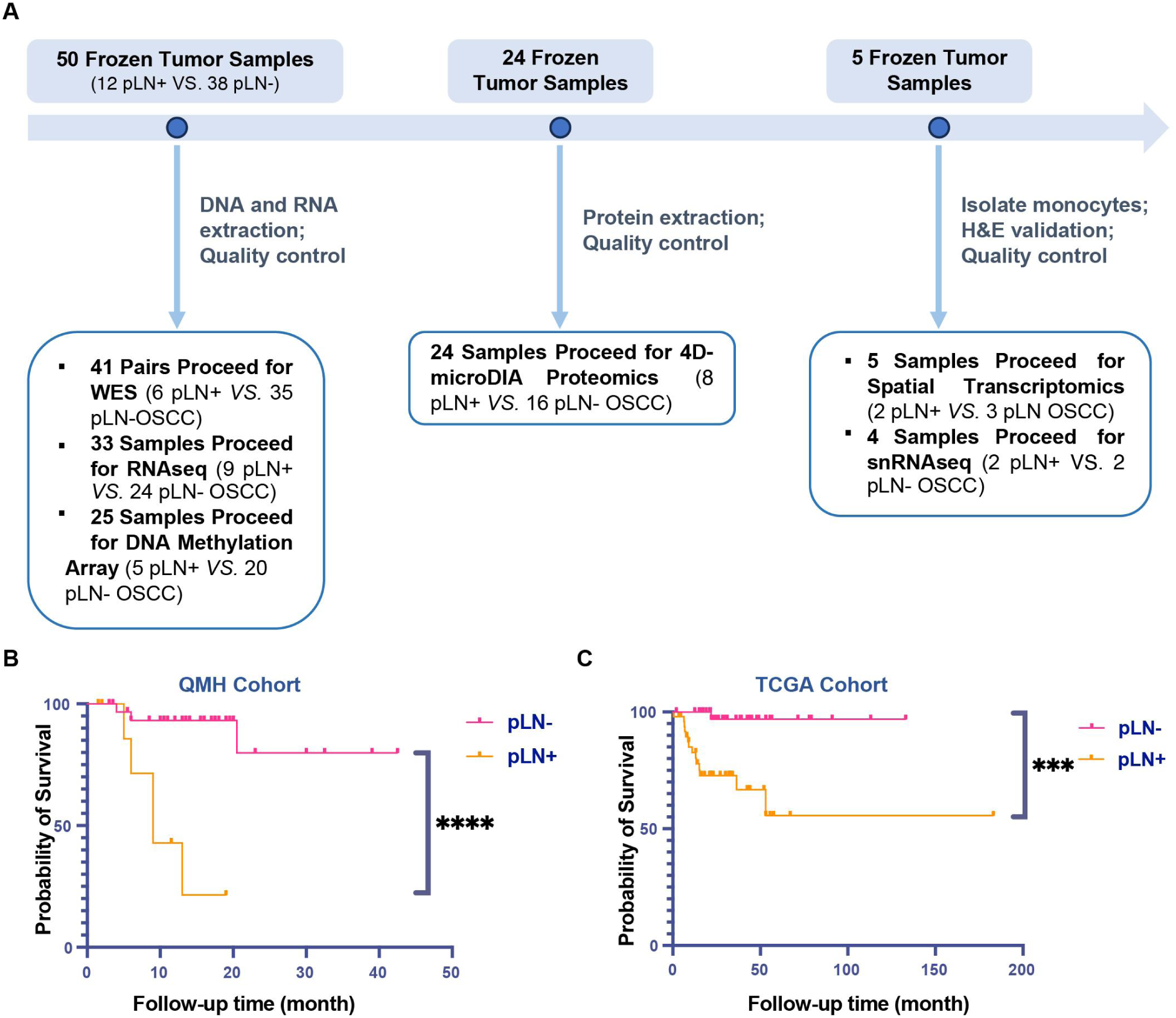
Experiment workflow and progression landscape of OSCC. **A.** Overview of sample processing workflow for this study. A cohort of 50 patients was initially included, from which a subset of 41 tumor samples and corresponding blood controls underwent whole-exome sequencing (WES). Additionally, 33 tumors were subjected to bulk tumor RNA sequencing, while 25 were analyzed using DNA methylation arrays. Protein extraction and quality control were performed on 24 fresh frozen tumor samples, which were subsequently subjected to 4D-micro DIA proteomics sequencing. Spatial transcriptomic analysis was performed on 5 fresh frozen samples, and single-nucleus RNA sequencing (snRNAseq) was performed on 4 samples. **B and C.** Kaplan–Meier survival curves comparing overall survival (OS) between OSCC patients with lymph node metastasis and those without lymph node metastasis in the QMH cohort **(B)** and the TCGA cohort **(C).**

**Supplementary Figure 2.**
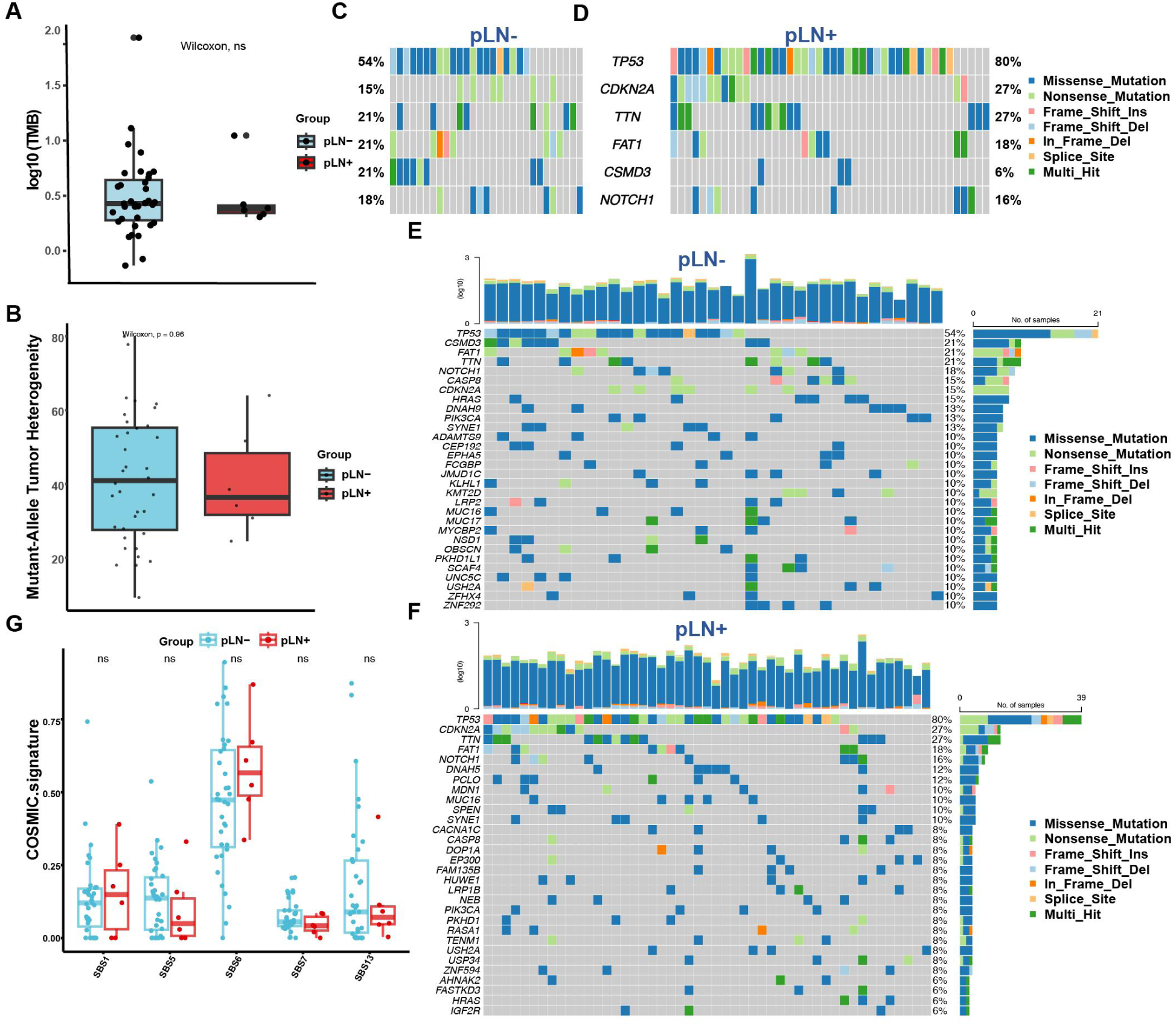
Supplemental gene mutation patterns. **A.** Box plots comparing mutational burden between primary OSCC with and without lymph node metastasis in the QMH cohort. P-value calculated using Wilcoxon tests. **B.** Box plots comparing mutant- allele tumor heterogeneity between primary OSCC with and without lymph node metastasis in the QMH cohort. P-value calculated using Wilcoxon tests. **C and D.** Co-Oncoplots showing shared and unique mutated genes in pLN- **(C)** and pLN+ **(D)** tumor samples in the TCGA cohort. **E and F.** Oncoplots showing individual mutated gene patterns in pLN- **(E)** and pLN+ **(F)** tumor samples in the TCGA cohort. **G.** Box plots comparing COSMIC signatures between primary OSCC with and without lymph node metastasis in the QMH cohort.

**Supplementary Figure 3.**
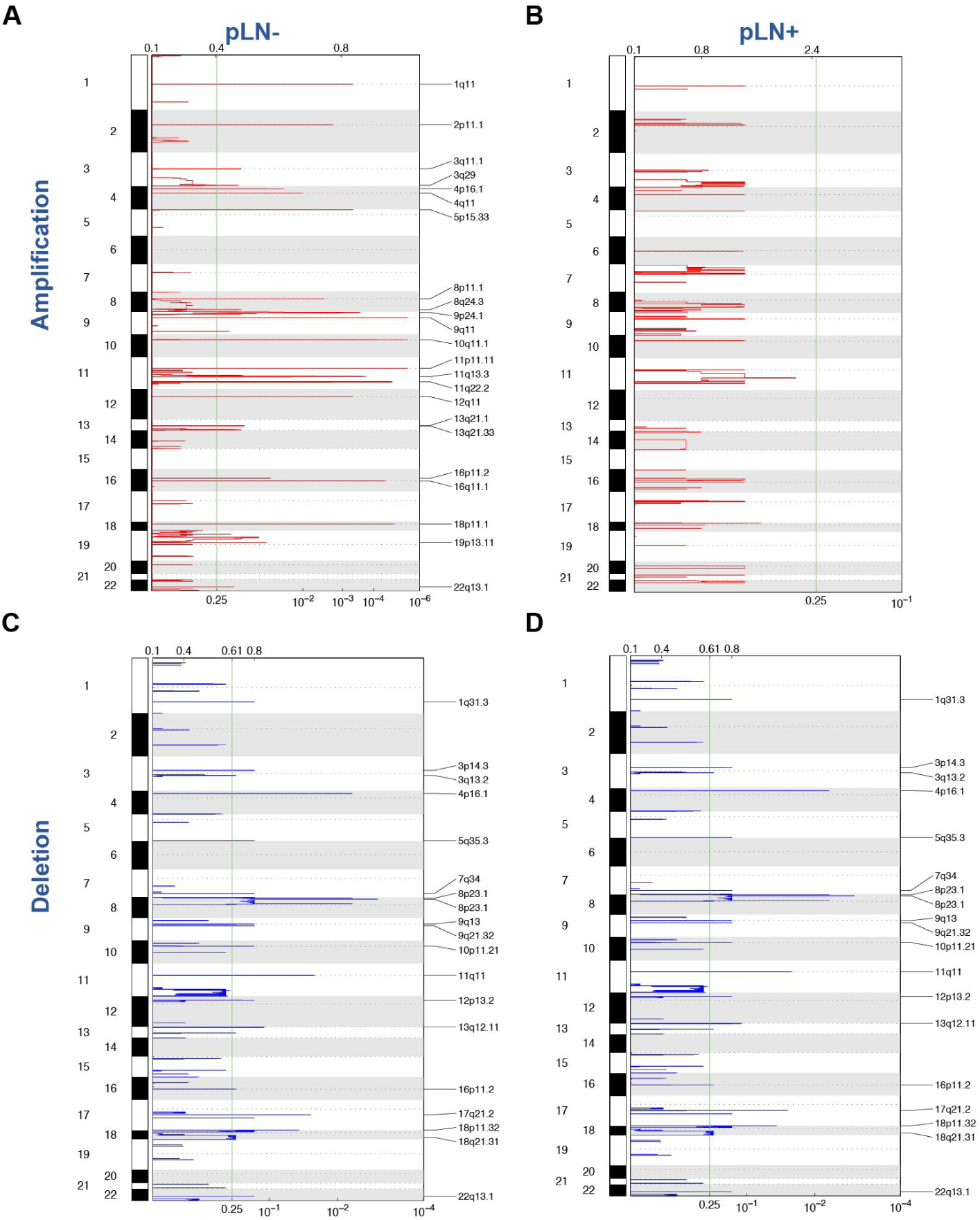
Landscape of somatic copy number alterations in pLN- and pLN+ tumor samples. Plots showing genomic locations and calculated q-values for aberrant regions as determined by the Genomic Identification of Significant Targets in Cancer (GISTIC) method for tumor samples in the QMH cohort. **A and C.** Amplification and depletion regions reported for pLN-. **B and D.** Amplification and depletion regions reported for pLN+. The genome is oriented vertically from top to bottom, with GISTIC scores at each locus plotted from left to right at the top and q-values plotted on a log scale at the bottom. The green line represents the significance threshold (q-value = 0.25).

**Supplementary Figure 4.**
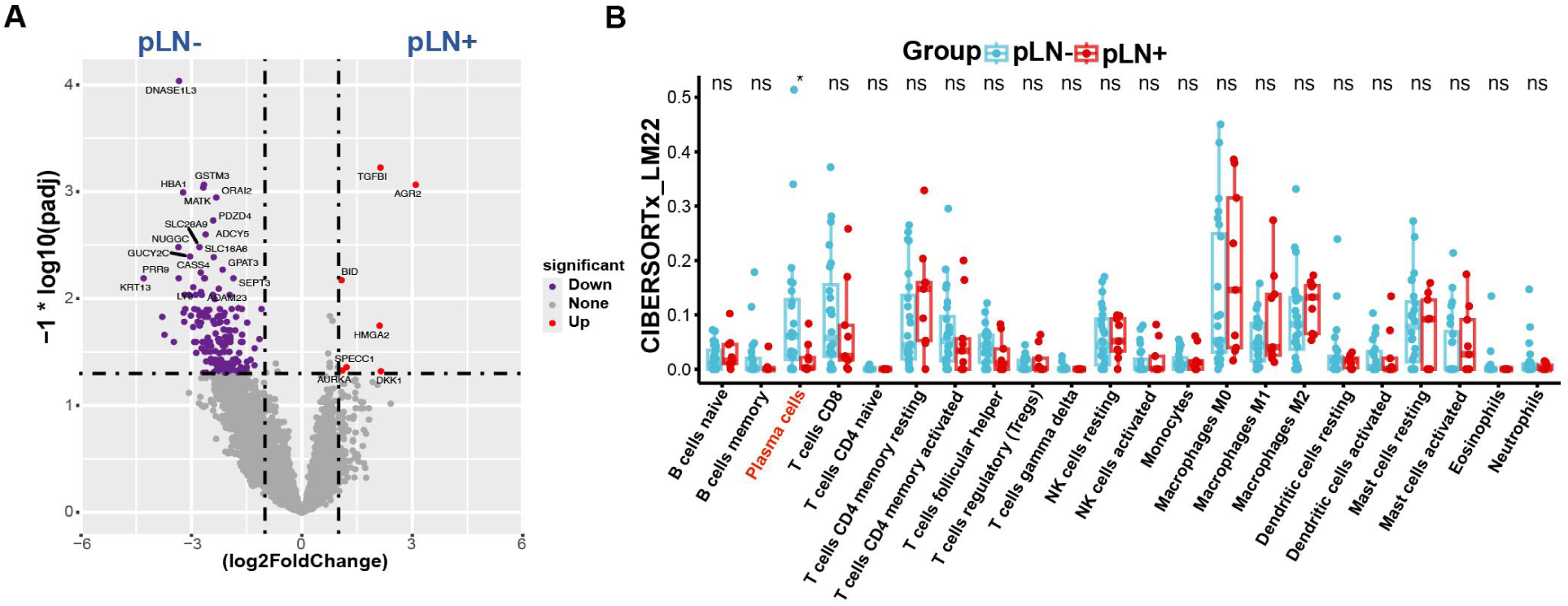
Differentially expressed genes (DEG) and CIBERSORTx findings from bulk tumor RNAseq. **A.** Volcano plot showing differentially expressed genes between pLN- and pLN+ groups. **B.** Box plot representing CIBERSORTx scores comparison between primary OSCC with and without lymph node metastasis.

**Supplementary Figure 5.**
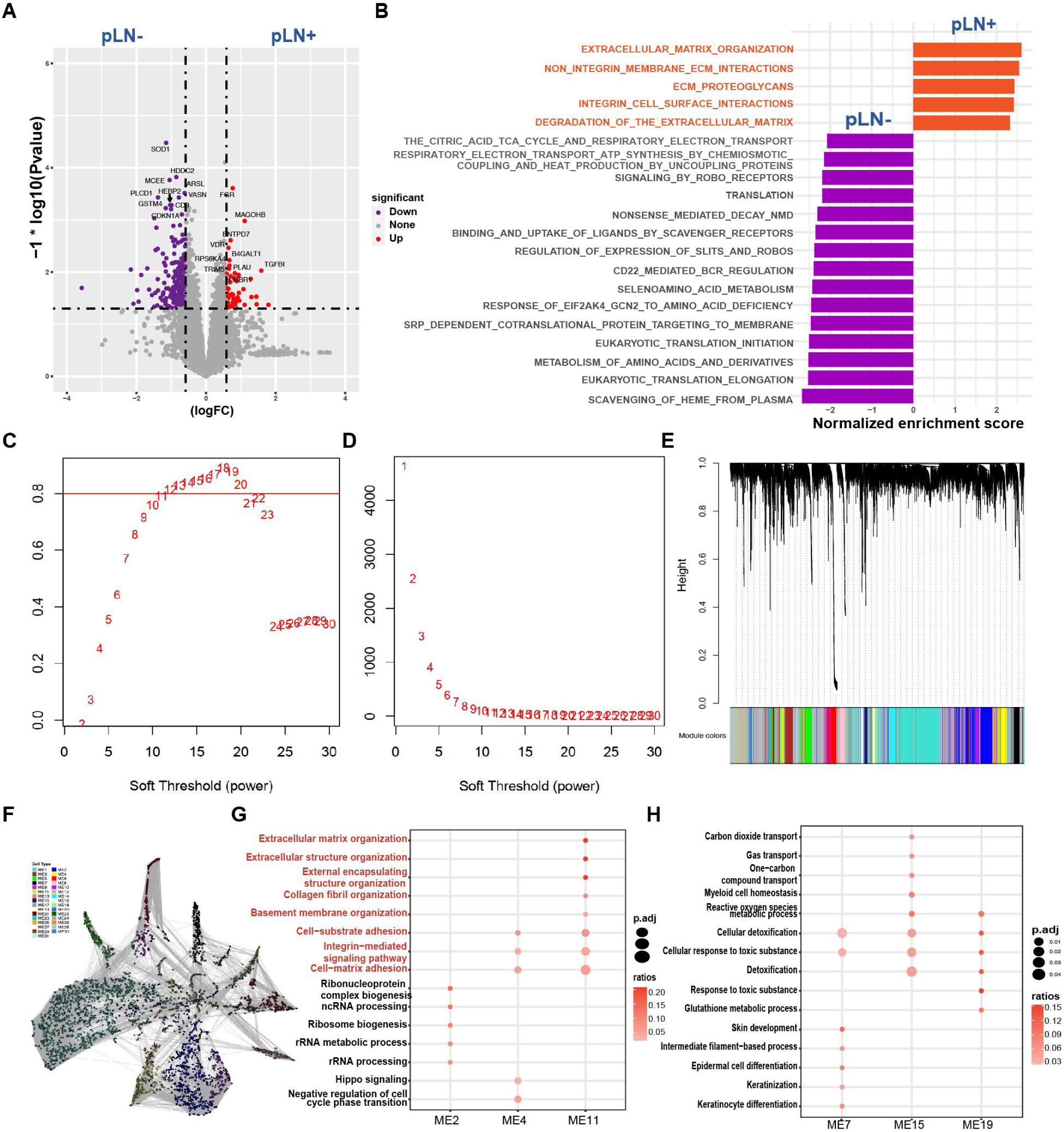
Proteomic analysis revealing differentially expressed proteins (DEP), GSEA, and construction of protein co-expression network. **A.** Volcano plot showing differentially expressed proteins between pLN- and pLN+ groups of 24 patients. **B.** Gene Set Enrichment Analysis (GSEA) plots of the top 20 pathways (ranked by adj. P values) based on the Reactome collection from MSigDB. The red bar towards the right indicates pathways enriched in primary OSCC with lymph node metastasis, and the purple bar towards the left indicates pathways enriched in primary OSCC without lymph node metastasis. Pathways enriched in pLN+ OSCC related to ECM remodeling are highlighted in red. **C and D.** Analysis of the scale-free fit index **(C)** and the mean connectivity **(D)** for determining soft-thresholding powers. **E.** Hierarchical clustering dendrogram of proteins in different modules. **F.** Protein co-expression network. Nodes are color-coded according to module membership. **G and H.** Functional enrichment in significantly dysregulated modules in the pLN+ group **(G)** and pLN- group **(H)**. Colors represent the relative ratios of significant genes within each pathway, circle size represents the adjusted p-values of each program in modules, and the x-axis represents the individual module number.

**Supplementary Figure 6.**
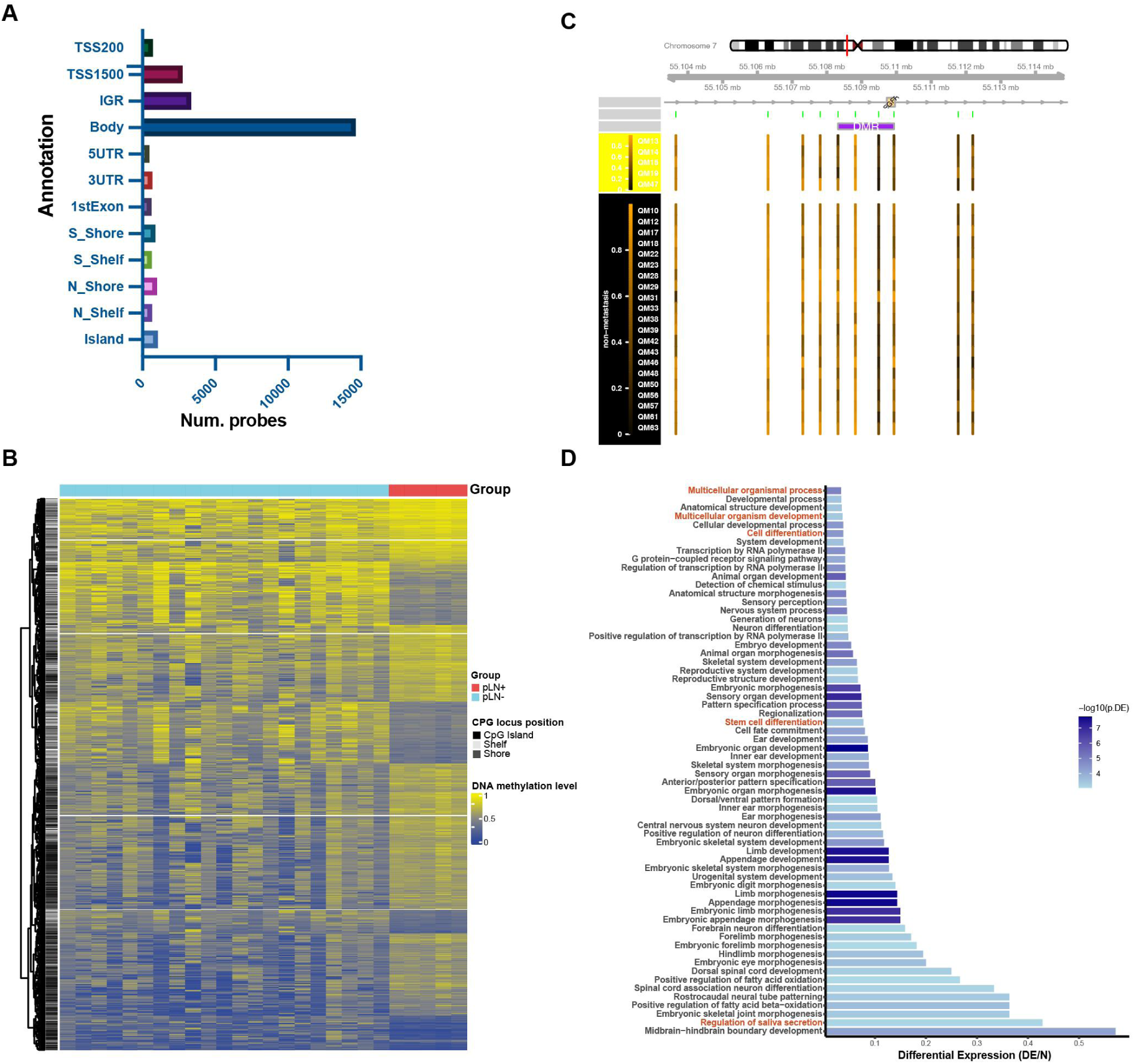
DNA methylation landscape of primary OSCC tumors. **A.** Distribution of differentially methylated probes (DMPs) in different regions related to CpG islands, including islands, CpG shores, and CpG shelves, and gene regions (TSS1500, TSS200, 5′ UTRs, first exons, gene bodies, and 3′ UTRs). **B.** Heatmap showing the methylation probe location of CpG islands, shores, and shelves. **C.** Differentially methylated regions covering the *EGFR* gene on chromosome 7. **D.** GO-BP pathway analysis of DMPs.

**Supplementary Figure 7.**
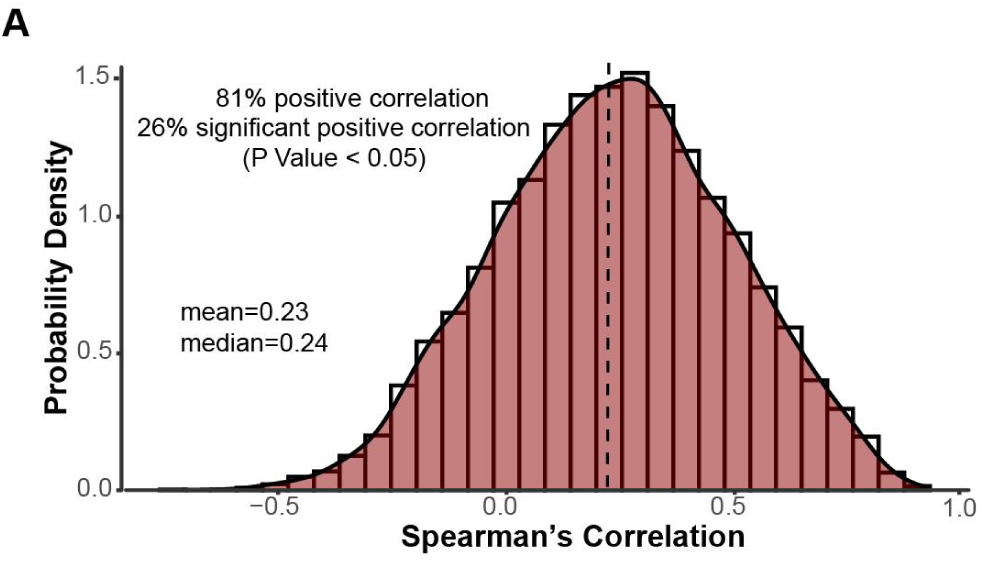
Distribution of Spearman’s correlation coefficients. **A.** Distribution of Spearman’s correlation coefficients between mRNA and protein log2-fold changes of individual genes across patients.

**Supplementary Figure 8.**
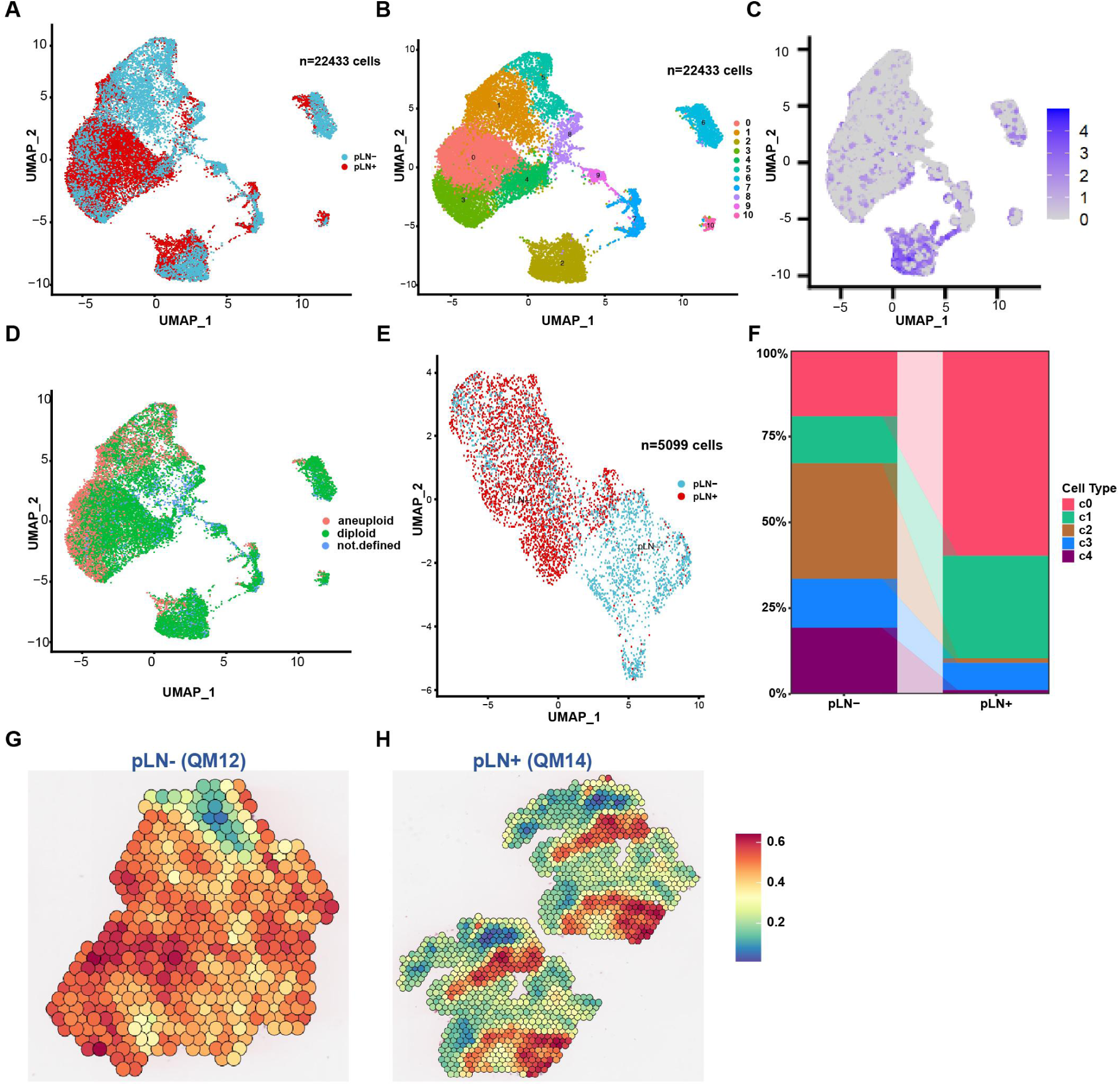
UMAP plot of cells distinguished by groups. **A.** UMAP plot of the single-cell profile colored by pLN+ and pLN- groups. **B.** UMAP plot of the single-cell profile colored by 10 clusters. **C.** UMAP atlas showing the expression levels of POSTN. **D.** UMAP plot showing CopyKat-screened cancer cells with extensive genome-wide copy number aberrations (aneuploidy). **E.** UMAP plot of the cancer-cell profile colored by pLN+ and pLN- groups. **F.** Cell abundance analysis showing the proportion of each cluster in pLN+ and pLN- OSCC. **G and H.** Spatial spots showing the enrichment level of EMT pathway in pLN- **(G)** and pLN+ OSCC **(H)**.

## Reference

1. Johnson NW, Jayasekara P, Amarasinghe AA. Squamous cell carcinoma and precursor lesions of the oral cavity: epidemiology and aetiology. Periodontol 2000. 2011;57(1):19–37.

2. Wang X, Xu J, Wang L, Liu C, Wang H. The role of cigarette smoking and alcohol consumption in the differentiation of oral squamous cell carcinoma for the males in China. J Cancer Res Ther. 2015;11(1):141–5.

3. Ram H, Sarkar J, Kumar H, Konwar R, Bhatt ML, Mohammad S. Oral cancer: risk factors and molecular pathogenesis. J Maxillofac Oral Surg. 2011;10(2):132–7.

4. Sharma A, Indu S, Gautami D, Sharma D. Oral squamous cell carcinoma (OSCC) in humans: Etiological Factors, diagnostic and therapeutic relevance. Research journal of biotechnology. 2020;15:141–51.

5. Meier JK, Schuderer JG, Zeman F, Klingelhöffer C, Hullmann M, Spanier G, et al. Health-related quality of life: a retrospective study on local vs. microvascular reconstruction in patients with oral cancer. BMC Oral Health. 2019;19(1):62.

6. González-Moles M, Aguilar-Ruiz M, Ramos-García P. Challenges in the Early Diagnosis of Oral Cancer, Evidence Gaps and Strategies for Improvement: A Scoping Review of Systematic Reviews. Cancers (Basel). 2022;14(19).

7. Chow LQM. Head and Neck Cancer. N Engl J Med. 2020;382(1):60–72.

8. Ho AS, Kim S, Tighiouart M, Gudino C, Mita A, Scher KS, et al. Metastatic Lymph Node Burden and Survival in Oral Cavity Cancer. J Clin Oncol. 2017;35(31):3601–9.

9. Siegel RL, Miller KD, Fuchs HE, Jemal A. Cancer statistics, 2021. Ca Cancer J Clin. 2021;71(1):7–33.

10. Shaikh S, Yadav DK, Bhadresha K, Rawal RM. Integrated computational screening and liquid biopsy approach to uncover the role of biomarkers for oral cancer lymph node metastasis. Scientific Reports. 2023;13(1):14033.

11. Ghantous Y, Mozalbat S, Nashef A, Abdol-Elraziq M, Sudri S, Araidy S, et al. EMT Dynamics in Lymph Node Metastasis of Oral Squamous Cell Carcinoma. Cancers. 2024;16(6):1185.

12. Thiery JP, Sleeman JP. Complex networks orchestrate epithelial-mesenchymal transitions. Nat Rev Mol Cell Biol. 2006;7(2):131–42.

13. Katsuno Y, Derynck R. Epithelial plasticity, epithelial-mesenchymal transition, and the TGF- &#x3b2; family. Developmental Cell. 2021;56(6):726–46.

14. Banerjee S, Lo W-C, Majumder P, Roy D, Ghorai M, Shaikh NK, et al. Multiple roles for basement membrane proteins in cancer progression and EMT. European Journal of Cell Biology. 2022;101(2):151220.

15. Sticht C, Hofele C, Flechtenmacher C, Bosch FX, Freier K, Lichter P, Joos S. Amplification of Cyclin L1 is associated with lymph node metastases in head and neck squamous cell carcinoma (HNSCC). Br J Cancer. 2005;92(4):770–4.

16. Dharavath B, Butle A, Pal A, Desai S, Upadhyay P, Rane A, et al. Role of miR-944/MMP10/AXL- axis in lymph node metastasis in tongue cancer. Communications Biology. 2023;6(1):57.

17. Liu S, Liu L, Ye W, Ye D, Wang T, Guo W, et al. High Vimentin Expression Associated with Lymph Node Metastasis and Predicated a Poor Prognosis in Oral Squamous Cell Carcinoma. Sci Rep. 2016;6:38834.

18. Shih CH, Chang YJ, Huang WC, Jang TH, Kung HJ, Wang WC, et al. EZH2-mediated upregulation of ROS1 oncogene promotes oral cancer metastasis. Oncogene. 2017;36(47):6542–54.

19. Hao Y, Xiao Y, Liao X, Tang S, Xie X, Liu R, Chen Q. FGF8 induces epithelial-mesenchymal transition and promotes metastasis in oral squamous cell carcinoma. Int J Oral Sci. 2021;13(1):6.

20. Horny K, Sproll C, Peiffer L, Furtmann F, Gerhardt P, Gravemeyer J, et al. Mesenchymal– epithelial transition in lymph node metastases of oral squamous cell carcinoma is accompanied by ZEB1 expression. Journal of Translational Medicine. 2023;21(1):267.

21. Ji X, Sun T, Xie S, Qian H, Song L, Wang L, et al. Upregulation of CPNE7 in mesenchymal stromal cells promotes oral squamous cell carcinoma metastasis through the NF-κB pathway. Cell Death Discov. 2021;7(1):294.

22. Cui B, Chen J, Luo M, Liu Y, Chen H, Lü D, et al. PKD3 promotes metastasis and growth of oral squamous cell carcinoma through positive feedback regulation with PD-L1 and activation of ERK- STAT1/3-EMT signalling. International Journal of Oral Science. 2021;13(1):8.

23. Pidugu VK, Wu MM, Yen AH, Pidugu HB, Chang KW, Liu CJ, Lee TC. IFIT1 and IFIT3 promote oral squamous cell carcinoma metastasis and contribute to the anti-tumor effect of gefitinib via enhancing p-EGFR recycling. Oncogene. 2019;38(17):3232–47.

24. Liu ZL, Meng XY, Bao RJ, Shen MY, Sun JJ, Chen WD, et al. Single cell deciphering of progression trajectories of the tumor ecosystem in head and neck cancer. Nature Communications. 2024;15(1):2595.

25. Choi J-H, Lee B-S, Jang JY, Lee YS, Kim HJ, Roh J, et al. Single-cell transcriptome profiling of the stepwise progression of head and neck cancer. Nature Communications. 2023;14(1):1055.

26. Eric H, Piersiala K, Lagebro V, Farrajota Neves Da Silva P, Petro M, Starkhammar M, et al. High expression of PD-L1 on conventional dendritic cells in tumour-draining lymph nodes is associated with poor prognosis in oral cancer. Cancer Immunology, Immunotherapy. 2024;73(9):165.

27. Rahim MK, Okholm TLH, Jones KB, McCarthy EE, Liu CC, Yee JL, et al. Dynamic CD8(+) T cell responses to cancer immunotherapy in human regional lymph nodes are disrupted in metastatic lymph nodes. Cell. 2023;186(6):1127–43.e18.

28. Melo-Alvim C, Neves ME, Santos JL, Abrunhosa-Branquinho AN, Barroso T, Costa L, Ribeiro L. Radiotherapy, Chemotherapy and Immunotherapy-Current Practice and Future Perspectives for Recurrent/Metastatic Oral Cavity Squamous Cell Carcinoma. Diagnostics (Basel). 2022;13(1).

29. Bhat GR, Sethi I, Sadida HQ, Rah B, Mir R, Algehainy N, et al. Cancer cell plasticity: from cellular, molecular, and genetic mechanisms to tumor heterogeneity and drug resistance. Cancer and Metastasis Reviews. 2024;43(1):197–228.

30. Pickering CR, Zhang J, Yoo SY, Bengtsson L, Moorthy S, Neskey DM, et al. Integrative genomic characterization of oral squamous cell carcinoma identifies frequent somatic drivers. Cancer Discov. 2013;3(7):770–81.

31. Gillison ML, Akagi K, Xiao W, Jiang B, Pickard RKL, Li J, et al. Human papillomavirus and the landscape of secondary genetic alterations in oral cancers. Genome Res. 2019;29(1):1–17.

32. Sequeira I, Rashid M, Tomás IM, Williams MJ, Graham TA, Adams DJ, et al. Genomic landscape and clonal architecture of mouse oral squamous cell carcinomas dictate tumour ecology. Nature Communications. 2020;11(1):5671.

33. Tan Y, Wang Z, Xu M, Li B, Huang Z, Qin S, et al. Oral squamous cell carcinomas: state of the field and emerging directions. Int J Oral Sci. 2023;15(1):44.

34. Shridhar K, Walia GK, Aggarwal A, Gulati S, Geetha AV, Prabhakaran D, et al. DNA methylation markers for oral pre-cancer progression: A critical review. Oral Oncol. 2016;53:1–9.

35. Elmusrati A, Wang J, Wang CY. Tumor microenvironment and immune evasion in head and neck squamous cell carcinoma. Int J Oral Sci. 2021;13(1):24.

36. Pour M, Yanai I. New adventures in spatial transcriptomics. Dev Cell. 2022;57(10):1209–10.

37. Wu D, Liu X, Zhang J, Li L, Wang X. Significance of single-cell and spatial transcriptomes in cell biology and toxicology. Cell Biology and Toxicology. 2021;37(1):1–5.

38. Ji H, Hu C, Yang X, Liu Y, Ji G, Ge S, et al. Lymph node metastasis in cancer progression: molecular mechanisms, clinical significance and therapeutic interventions. Signal Transduction and Targeted Therapy. 2023;8(1):367.

39. Comprehensive genomic characterization of head and neck squamous cell carcinomas. Nature. 2015;517(7536):576–82.

40. Benjamin D, Sato T, Cibulskis K, Getz G, Stewart C, Lichtenstein L. Calling Somatic SNVs and Indels with Mutect2. bioRxiv. 2019:861054.

41. Sha D, Jin Z, Budczies J, Kluck K, Stenzinger A, Sinicrope FA. Tumor Mutational Burden as a Predictive Biomarker in Solid Tumors. Cancer Discov. 2020;10(12):1808–25.

42. Mroz EA, Rocco JW. MATH, a novel measure of intratumor genetic heterogeneity, is high in poor- outcome classes of head and neck squamous cell carcinoma. Oral oncology. 2013;49(3):211–5.

43. Zhao Q, Zheng H, Duan W, Li C, Xie W, Wang G, et al. Association between MUC5B mutation and prognosis across solid tumors. Journal of Clinical Oncology. 2020;38(15_suppl):e13515–e.

44. Jiang X, He Y, Shen Q, Duan L, Yuan Y, Tang L, et al. RETSAT Mutation Selected for Hypoxia Adaptation Inhibits Tumor Growth. Front Cell Dev Biol. 2021;9:744992.

45. Wang H, Guo M, Wei H, Chen Y. Targeting p53 pathways: mechanisms, structures, and advances in therapy. Signal Transduction and Targeted Therapy. 2023;8(1):92.

46. Liang J, Fan J, Wang M, Niu Z, Zhang Z, Yuan L, et al. CDKN2A inhibits formation of homotypic cell-in-cell structures. Oncogenesis. 2018;7(6):50.

47. Sondka Z, Dhir NB, Carvalho-Silva D, Jupe S, Madhumita, McLaren K, et al. COSMIC: a curated database of somatic variants and clinical data for cancer. Nucleic Acids Research. 2023;52(D1):D1210–D7.

48. Li X, He Y, Wu J, Qiu J, Li J, Wang Q, et al. A novel pathway mutation perturbation score predicts the clinical outcomes of immunotherapy. Briefings in Bioinformatics. 2022;23(5).

49. Alcami A. Viral mimicry of cytokines, chemokines and their receptors. Nature Reviews Immunology. 2003;3(1):36–50.

50. Mermel CH, Schumacher SE, Hill B, Meyerson ML, Beroukhim R, Getz G. GISTIC2.0 facilitates sensitive and confident localization of the targets of focal somatic copy-number alteration in human cancers. Genome Biology. 2011;12(4):R41.

51. Chang X, Zhao Y, Hou C, Glessner J, McDaniel L, Diamond MA, et al. Common variants in MMP20 at 11q22.2 predispose to 11q deletion and neuroblastoma risk. Nat Commun. 2017;8(1):569.

52. Dubard Gault M, Mandelker D, DeLair D, Stewart CR, Kemel Y, Sheehan MR, et al. Germline SDHA mutations in children and adults with cancer. Cold Spring Harb Mol Case Stud. 2018;4(4).

53. Li J, Peng Y. Knockdown RPL29 Gene Can Inhibit the Proliferation, Invasion of Squamous Cell Carcinomas. Ann Clin Lab Sci. 2019;49(6):763–9.

54. Shi M, Huang K, Wei J, Wang S, Yang W, Wang H, Li Y. Identification and Validation of a Prognostic Signature Derived from the Cancer Stem Cells for Oral Squamous Cell Carcinoma. Int J Mol Sci. 2024;25(2).

55. Xie J-M, Li B, Yu H-P, Gao Q-G, Li W, Wu H-R, Qin Z-H. TIGAR Has a Dual Role in Cancer Cell Survival through Regulating Apoptosis and Autophagy. Cancer Research. 2014;74(18):5127–38.

56. Li Y, Wan Q, Wang W, Mai L, Sha L, Mashrah M, et al. LncRNA ADAMTS9-AS2 promotes tongue squamous cell carcinoma proliferation, migration and EMT via the miR-600/EZH2 axis. Biomedicine & Pharmacotherapy. 2019;112:108719.

57. Bruckman KC, Schönleben F, Qiu W, Woo VL, Su GH. Mutational analyses of the BRAF, KRAS, and PIK3CA genes in oral squamous cell carcinoma. Oral Surg Oral Med Oral Pathol Oral Radiol Endod. 2010;110(5):632–7.

58. Hartanto FK, Karen-Ng LP, Vincent-Chong VK, Ismail SM, Mustafa WM, Abraham MT, et al. KRT13, FAIM2 and CYP2W1 mRNA expression in oral squamous cell carcinoma patients with risk habits. Asian Pac J Cancer Prev. 2015;16(3):953–8.

59. Guan C, Ouyang D, Qiao Y, Li K, Zheng G, Lao X, et al. CA9 transcriptional expression determines prognosis and tumour grade in tongue squamous cell carcinoma patients. Journal of Cellular and Molecular Medicine. 2020;24(10):5832–41.

60. Mao L, Lee JS, Fan YH, Ro JY, Batsakis JG, Lippman S, et al. Frequent microsatellite alterations at chromosomes 9p21 and 3p14 in oral premalignant lesions and their value in cancer risk assessment. Nature Medicine. 1996;2(6):682–5.

61. Bhosale PG, Pandey M, Cristea S, Shah M, Patil A, Beerenwinkel N, et al. Recurring Amplification at 11q22.1-q22.2 Locus Plays an Important Role in Lymph Node Metastasis and Radioresistance in OSCC. Sci Rep. 2017;7(1):16051.

62. Dong A, Wodziak D, Lowe AW. Epidermal Growth Factor Receptor (EGFR) Signaling Requires a Specific Endoplasmic Reticulum Thioredoxin for the Post-translational Control of Receptor Presentation to the Cell Surface*. Journal of Biological Chemistry. 2015;290(13):8016–27.

63. Wodziak D, Dong A, Basin MF, Lowe AW. Anterior Gradient 2 (AGR2) Induced Epidermal Growth Factor Receptor (EGFR) Signaling Is Essential for Murine Pancreatitis-Associated Tissue Regeneration. PLoS One. 2016;11(10):e0164968.

64. Derynck R, Turley SJ, Akhurst RJ. TGFβ biology in cancer progression and immunotherapy. Nat Rev Clin Oncol. 2021;18(1):9–34.

65. Xu J, Lamouille S, Derynck R. TGF-beta-induced epithelial to mesenchymal transition. Cell Res. 2009;19(2):156–72.

66. Mou PK, Yang EJ, Shi C, Ren G, Tao S, Shim JS. Aurora kinase A, a synthetic lethal target for precision cancer medicine. Experimental & Molecular Medicine. 2021;53(5):835–47.

67. Liberzon A, Birger C, Thorvaldsdóttir H, Ghandi M, Mesirov JP, Tamayo P. The Molecular Signatures Database (MSigDB) hallmark gene set collection. Cell Syst. 2015;1(6):417–25.

68. Garofano L, Migliozzi S, Oh YT, D’Angelo F, Najac RD, Ko A, et al. Pathway-based classification of glioblastoma uncovers a mitochondrial subtype with therapeutic vulnerabilities. Nature Cancer. 2021;2(2):141–56.

69. Subramanian A, Tamayo P, Mootha VK, Mukherjee S, Ebert BL, Gillette MA, et al. Gene set enrichment analysis: A knowledge-based approach for interpreting genome-wide expression profiles. Proceedings of the National Academy of Sciences. 2005;102(43):15545–50.

70. Löbrich M, Jeggo PA. The impact of a negligent G2/M checkpoint on genomic instability and cancer induction. Nature Reviews Cancer. 2007;7(11):861–9.

71. Segeren HA, van Rijnberk LM, Moreno E, Riemers FM, van Liere EA, Yuan R, et al. Excessive E2F Transcription in Single Cancer Cells Precludes Transient Cell-Cycle Exit after DNA Damage. Cell Reports. 2020;33(9).

72. Kerseviciute I, Gordevicius J. aPEAR: an R package for autonomous visualization of pathway enrichment networks. Bioinformatics. 2023;39(11).

73. Chen B, Khodadoust MS, Liu CL, Newman AM, Alizadeh AA. Profiling Tumor Infiltrating Immune Cells with CIBERSORT. Methods Mol Biol. 2018;1711:243–59.

74. Yoshihara K, Shahmoradgoli M, Martínez E, Vegesna R, Kim H, Torres-Garcia W, et al. Inferring tumour purity and stromal and immune cell admixture from expression data. Nature Communications. 2013;4(1):2612.

75. Aran D, Hu Z, Butte AJ. xCell: digitally portraying the tissue cellular heterogeneity landscape. Genome Biology. 2017;18(1):220.

76. Fraga M, Yáñez M, Sherman M, Llerena F, Hernandez M, Nourdin G, et al. Immunomodulation of T Helper Cells by Tumor Microenvironment in Oral Cancer Is Associated With CCR8 Expression and Rapid Membrane Vitamin D Signaling Pathway. Front Immunol. 2021;12:643298.

77. Feng Q, Wei H, Morihara J, Stern J, Yu M, Kiviat N, et al. Th2 type inflammation promotes the gradual progression of HPV-infected cervical cells to cervical carcinoma. Gynecologic Oncology. 2012;127(2):412–9.

78. Alam A, Levanduski E, Denz P, Villavicencio HS, Bhatta M, Alhorebi L, et al. Fungal mycobiome drives IL-33 secretion and type 2 immunity in pancreatic cancer. Cancer Cell. 2022;40(2):153–67.e11.

79. Fässler M, Diem S, Mangana J, Hasan Ali O, Berner F, Bomze D, et al. Antibodies as biomarker candidates for response and survival to checkpoint inhibitors in melanoma patients. J Immunother Cancer. 2019;7(1):50.

80. Chen Y, Sun J, Luo Y, Liu J, Wang X, Feng R, et al. Pharmaceutical targeting Th2-mediated immunity enhances immunotherapy response in breast cancer. Journal of Translational Medicine. 2022;20(1):615.

81. Gillet LC, Navarro P, Tate S, Röst H, Selevsek N, Reiter L, et al. Targeted data extraction of the MS/MS spectra generated by data-independent acquisition: a new concept for consistent and accurate proteome analysis. Mol Cell Proteomics. 2012;11(6):O111.016717.

82. Qiu M, Lin Q, Liu Y, Chen P, Zhou Y, Jiang Y, et al. Potentially functional genetic variants in RPS6KA4 and MAP2K5 in the MAPK signaling pathway predict HBV-related hepatocellular carcinoma survival. Mol Carcinog. 2023;62(9):1378–87.

83. Chen G, Sun J, Xie M, Yu S, Tang Q, Chen L. PLAU Promotes Cell Proliferation and Epithelial- Mesenchymal Transition in Head and Neck Squamous Cell Carcinoma. Frontiers in Genetics. 2021;12.

84. Szalmás A, Tomaić V, Basukala O, Massimi P, Mittal S, Kónya J, Banks L. The PTPN14 Tumor Suppressor Is a Degradation Target of Human Papillomavirus E7. Journal of Virology. 2017;91(7):10.1128/jvi.00057-17.

85. Yuan Z, Li Y, Zhang S, Wang X, Dou H, Yu X, et al. Extracellular matrix remodeling in tumor progression and immune escape: from mechanisms to treatments. Molecular Cancer. 2023;22(1):48.

86. Langfelder P, Horvath S. WGCNA: an R package for weighted correlation network analysis. BMC Bioinformatics. 2008;9(1):559.

87. AmeliMojarad M, AmeliMojarad M, Cui X, shariati P. Pan-cancer analysis of CTNNB1 with potential as a therapeutic target for human tumorigenesis. Informatics in Medicine Unlocked. 2023;42:101331.

88. van Schie EH, van Amerongen R. Aberrant WNT/CTNNB1 Signaling as a Therapeutic Target in Human Breast Cancer: Weighing the Evidence. Front Cell Dev Biol. 2020;8:25.

89. Thomas de Montpréville V, Lacroix L, Rouleau E, Mamodaly M, Leclerc J, Tutuianu L, et al. Non- small cell lung carcinomas with CTNNB1 (beta-catenin) mutations: A clinicopathological study of 26 cases. Annals of Diagnostic Pathology. 2020;46:151522.

90. Hema KN, Smitha T, Sheethal HS, Mirnalini SA. Epigenetics in oral squamous cell carcinoma. J Oral Maxillofac Pathol. 2017;21(2):252–9.

91. Goldberg AD, Allis CD, Bernstein E. Epigenetics: a landscape takes shape. Cell. 2007;128(4):635–8.

92. Basu B, Chakraborty J, Chandra A, Katarkar A, Baldevbhai JRK, Dhar Chowdhury D, et al. Genome-wide DNA methylation profile identified a unique set of differentially methylated immune genes in oral squamous cell carcinoma patients in India. Clin Epigenetics. 2017;9:13.

93. Xu Z, Qin F, Yuan L, Wei J, Sun Y, Qin J, et al. EGFR DNA Methylation Correlates With EGFR Expression, Immune Cell Infiltration, and Overall Survival in Lung Adenocarcinoma. Front Oncol. 2021;11:691915.

94. Liu AY, Zheng H, Ouyang G. Periostin, a multifunctional matricellular protein in inflammatory and tumor microenvironments. Matrix Biology. 2014;37:150–6.

95. Bessa X, Elizalde JI, Mitjans F, Piñol V, Miquel R, Panés J, et al. Leukocyte recruitment in colon cancer: role of cell adhesion molecules, nitric oxide, and transforming growth factor beta1. Gastroenterology. 2002;122(4):1122–32.

96. Liao J, Chen R, Lin B, Deng R, Liang Y, Zeng J, et al. Cross-Talk between the TGF-β and Cell Adhesion Signaling Pathways in Cancer. Int J Med Sci. 2024;21(7):1307–20.

97. Meacham CE, Morrison SJ. Tumour heterogeneity and cancer cell plasticity. Nature. 2013;501(7467):328–37.

98. Weinberg RA. Coming full circle-from endless complexity to simplicity and back again. Cell. 2014;157(1):267–71.

99. Navin NE. The first five years of single-cell cancer genomics and beyond. Genome Res. 2015;25(10):1499–507.

100. Tanay A, Regev A. Scaling single-cell genomics from phenomenology to mechanism. Nature. 2017;541(7637):331-8.

101. Gao R, Bai S, Henderson YC, Lin Y, Schalck A, Yan Y, et al. Delineating copy number and clonal substructure in human tumors from single-cell transcriptomes. Nat Biotechnol. 2021;39(5):599–608.

102. Wright K, Ly T, Kriet M, Czirok A, Thomas SM. Cancer-Associated Fibroblasts: Master Tumor Microenvironment Modifiers. Cancers (Basel). 2023;15(6).

103. Li X, Wang C-Y. From bulk, single-cell to spatial RNA sequencing. International Journal of Oral Science. 2021;13(1):36.

104. Williams CG, Lee HJ, Asatsuma T, Vento-Tormo R, Haque A. An introduction to spatial transcriptomics for biomedical research. Genome Medicine. 2022;14(1):68.

105. Oral cancer - the fight must go on against all odds…. Evidence-Based Dentistry. 2022;23(1):4–5.

106. Parmar A, Macluskey M, Mc Goldrick N, Conway DI, Glenny AM, Clarkson JE, et al. Interventions for the treatment of oral cavity and oropharyngeal cancer: chemotherapy. Cochrane Database Syst Rev. 2021;12(12):Cd006386.

107. Gamez ME, Kraus R, Hinni ML, Moore EJ, Ma DJ, Ko SJ, et al. Treatment outcomes of squamous cell carcinoma of the oral cavity in young adults. Oral Oncol. 2018;87:43–8.

108. Lee NY, Ferris RL, Psyrri A, Haddad RI, Tahara M, Bourhis J, et al. Avelumab plus standard-of- care chemoradiotherapy versus chemoradiotherapy alone in patients with locally advanced squamous cell carcinoma of the head and neck: a randomised, double-blind, placebo-controlled, multicentre, phase 3 trial. Lancet Oncol. 2021;22(4):450–62.

109. Vogel C, Marcotte EM. Insights into the regulation of protein abundance from proteomic and transcriptomic analyses. Nature Reviews Genetics. 2012;13(4):227–32.

110. Chen YJ, Roumeliotis TI, Chang YH, Chen CT, Han CL, Lin MH, et al. Proteogenomics of Non- smoking Lung Cancer in East Asia Delineates Molecular Signatures of Pathogenesis and Progression. Cell. 2020;182(1):226–44.e17.

111. Altelaar AF, Munoz J, Heck AJ. Next-generation proteomics: towards an integrative view of proteome dynamics. Nat Rev Genet. 2013;14(1):35–48.

112. Hanash S. Disease proteomics. Nature. 2003;422(6928):226–32.

113. Mani DR, Krug K, Zhang B, Satpathy S, Clauser KR, Ding L, et al. Cancer proteogenomics: current impact and future prospects. Nat Rev Cancer. 2022;22(5):298–313.

114. Rodriguez H, Zenklusen JC, Staudt LM, Doroshow JH, Lowy DR. The next horizon in precision oncology: Proteogenomics to inform cancer diagnosis and treatment. Cell. 2021;184(7):1661–70.

115. Popova NV, Jücker M. The Functional Role of Extracellular Matrix Proteins in Cancer. Cancers (Basel). 2022;14(1).

116. Chakravarthy A, Khan L, Bensler NP, Bose P, De Carvalho DD. TGF-β-associated extracellular matrix genes link cancer-associated fibroblasts to immune evasion and immunotherapy failure. Nature Communications. 2018;9(1):4692.

117. Hinz B. The extracellular matrix and transforming growth factor-β1: Tale of a strained relationship. Matrix Biology. 2015;47:54–65.

118. Wang BJ, Chi KP, Shen RL, Zheng SW, Guo Y, Li JF, et al. TGFBI Promotes Tumor Growth and is Associated with Poor Prognosis in Oral Squamous Cell Carcinoma. J Cancer. 2019;10(20):4902–12.

119. Ahmed AA, Mills AD, Ibrahim AE, Temple J, Blenkiron C, Vias M, et al. The extracellular matrix protein TGFBI induces microtubule stabilization and sensitizes ovarian cancers to paclitaxel. Cancer Cell. 2007;12(6):514–27.

120. Yokobori T, Nishiyama M. TGF-β Signaling in Gastrointestinal Cancers: Progress in Basic and Clinical Research. J Clin Med. 2017;6(1).

121. Reticker-Flynn NE, Zhang W, Belk JA, Basto PA, Escalante NK, Pilarowski GOW, et al. Lymph node colonization induces tumor-immune tolerance to promote distant metastasis. Cell. 2022;185(11):1924–42.e23.

122. Peng JM, Su YL. Lymph node metastasis and tumor-educated immune tolerance: Potential therapeutic targets against distant metastasis. Biochem Pharmacol. 2023;215:115731.

123. Liu Z, Zhang Z, Zhang Y, Zhou W, Zhang X, Peng C, et al. Spatial transcriptomics reveals that metabolic characteristics define the tumor immunosuppression microenvironment via iCAF transformation in oral squamous cell carcinoma. International Journal of Oral Science. 2024;16(1):9.

124. Hanahan D, Weinberg RA. The Hallmarks of Cancer. Cell. 2000;100(1):57–70.

125. Miao D, Margolis CA, Vokes NI, Liu D, Taylor-Weiner A, Wankowicz SM, et al. Genomic correlates of response to immune checkpoint blockade in microsatellite-stable solid tumors. Nature Genetics. 2018;50(9):1271–81.

126. Suski JM, Braun M, Strmiska V, Sicinski P. Targeting cell-cycle machinery in cancer. Cancer Cell. 2021;39(6):759–78.

127. Castro-Gamero AM, Pezuk JA, Brassesco MS, Tone LG. G2/M inhibitors as pharmacotherapeutic opportunities for glioblastoma: the old, the new, and the future. Cancer Biol Med. 2018;15(4):354–74.

128. Dorafshan S, Razmi M, Safaei S, Gentilin E, Madjd Z, Ghods R. Periostin: biology and function in cancer. Cancer Cell International. 2022;22(1):315.

129. Xu C, Wang Z, Zhang L, Feng Y, Lv J, Wu Z, et al. Periostin promotes the proliferation and metastasis of osteosarcoma by increasing cell survival and activates the PI3K/Akt pathway. Cancer Cell International. 2022;22(1):34.

130. Chen L, Tian X, Gong W, Sun B, Li G, Liu D, et al. Periostin mediates epithelial-mesenchymal transition through the MAPK/ERK pathway in hepatoblastoma. Cancer Biol Med. 2019;16(1):89–100.

131. Chen Y, Zhang F, Zhang B, Trojanowicz B, Hämmerle M, Kleeff J, Sunami Y. Periostin is associated with prognosis and immune cell infiltration in pancreatic adenocarcinoma based on integrated bioinformatics analysis. Cancer Reports. 2024;7(2):e1990.

132. Gao F, Liu J, Gan H. The expression of POSTN and immune cell infiltration are prognostic factors of lung adenocarcinoma. Medicine (Baltimore). 2022;101(34):e30187.

133. Hu C, Zhang Y, Wu C, Huang Q. Heterogeneity of cancer-associated fibroblasts in head and neck squamous cell carcinoma: opportunities and challenges. Cell Death Discovery. 2023;9(1):124.

134. Li X, González-Maroto C, Tavassoli M. Crosstalk between CAFs and tumour cells in head and neck cancer. Cell Death Discovery. 2024;10(1):303.

135. Chen C, Guo Q, Liu Y, Hou Q, Liao M, Guo Y, et al. Single-cell and spatial transcriptomics reveal POSTN+ cancer-associated fibroblasts correlated with immune suppression and tumour progression in non-small cell lung cancer. Clinical and Translational Medicine. 2023;13(12):e1515.

136. Chalmers ZR, Connelly CF, Fabrizio D, Gay L, Ali SM, Ennis R, et al. Analysis of 100,000 human cancer genomes reveals the landscape of tumor mutational burden. Genome Medicine. 2017;9(1):34.

137. Talevich E, Shain AH, Botton T, Bastian BC. CNVkit: Genome-Wide Copy Number Detection and Visualization from Targeted DNA Sequencing. PLOS Computational Biology. 2016;12(4):e1004873.

138. Pertea M, Kim D, Pertea GM, Leek JT, Salzberg SL. Transcript-level expression analysis of RNA- seq experiments with HISAT, StringTie and Ballgown. Nature Protocols. 2016;11(9):1650–67.

139. Yu G, Wang LG, Han Y, He QY. clusterProfiler: an R package for comparing biological themes among gene clusters. Omics. 2012;16(5):284–7.

140. Jiao X, Sherman BT, Huang da W, Stephens R, Baseler MW, Lane HC, Lempicki RA. DAVID-WS: a stateful web service to facilitate gene/protein list analysis. Bioinformatics. 2012;28(13):1805–6.

141. Szklarczyk D, Gable AL, Nastou KC, Lyon D, Kirsch R, Pyysalo S, et al. The STRING database in 2021: customizable protein-protein networks, and functional characterization of user-uploaded gene/measurement sets. Nucleic Acids Res. 2021;49(D1):D605–d12.

142. Shannon P, Markiel A, Ozier O, Baliga NS, Wang JT, Ramage D, et al. Cytoscape: a software environment for integrated models of biomolecular interaction networks. Genome Res. 2003;13(11):2498–504.

143. Tian Y, Morris TJ, Webster AP, Yang Z, Beck S, Feber A, Teschendorff AE. ChAMP: updated methylation analysis pipeline for Illumina BeadChips. Bioinformatics. 2017;33(24):3982–4.

144. Phipson B, Maksimovic J, Oshlack A. missMethyl: an R package for analyzing data from Illumina’s HumanMethylation450 platform. Bioinformatics. 2016;32(2):286–8.

145. Park J, Kim J, Kim E, Kim WJ, Won S. Prenatal lead exposure and cord blood DNA methylation in the Korean Exposome Study. Environ Res. 2021;195:110767.

146. Liao Y, Wang J, Jaehnig EJ, Shi Z, Zhang B. WebGestalt 2019: gene set analysis toolkit with revamped UIs and APIs. Nucleic Acids Res. 2019;47(W1):W199–w205.

147. Alkaslasi MR, Piccus ZE, Hareendran S, Silberberg H, Chen L, Zhang Y, et al. Single nucleus RNA-sequencing defines unexpected diversity of cholinergic neuron types in the adult mouse spinal cord. Nature Communications. 2021;12(1):2471.

148. Le DT, Durham JN, Smith KN, Wang H, Bartlett BR, Aulakh LK, et al. Mismatch repair deficiency predicts response of solid tumors to PD-1 blockade. Science. 2017;357(6349):409-13.

149. Satija R, Farrell JA, Gennert D, Schier AF, Regev A. Spatial reconstruction of single-cell gene expression data. Nat Biotechnol. 2015;33(5):495–502.

150. Stuart T, Butler A, Hoffman P, Hafemeister C, Papalexi E, Mauck WM, 3rd, et al. Comprehensive Integration of Single-Cell Data. Cell. 2019;177(7):1888–902.e21.

151. Hu C, Li T, Xu Y, Zhang X, Li F, Bai J, et al. CellMarker 2.0: an updated database of manually curated cell markers in human/mouse and web tools based on scRNA-seq data. Nucleic Acids Research. 2022;51(D1):D870–D6.

152. Jin S, Guerrero-Juarez CF, Zhang L, Chang I, Ramos R, Kuan C-H, et al. Inference and analysis of cell-cell communication using CellChat. Nature Communications. 2021;12(1):1088.

153. Hänzelmann S, Castelo R, Guinney J. GSVA: gene set variation analysis for microarray and RNA-Seq data. BMC Bioinformatics. 2013;14(1):7.

